# Trans-synaptic molecular context of NMDA receptor nanodomains

**DOI:** 10.1101/2023.12.22.573055

**Authors:** Michael C Anderson, Poorna A Dharmasri, Martina Damenti, Sarah R Metzbower, Rozita Laghaei, Thomas A Blanpied, Aaron D Levy

## Abstract

Tight coordination of the spatial relationships between protein complexes is required for cellular function. In neuronal synapses, many proteins responsible for neurotransmission organize into subsynaptic nanoclusters whose trans-cellular alignment modulates synaptic signal propagation. However, the spatial relationships between these proteins and NMDA receptors (NMDARs), which are required for learning and memory, remain undefined. Here, we mapped the relationship of key NMDAR subunits to reference proteins in the active zone and postsynaptic density using multiplexed super-resolution DNA-PAINT microscopy. GluN2A and GluN2B subunits formed nanoclusters with diverse configurations that, surprisingly, were not localized near presynaptic vesicle release sites marked by Munc13-1. Despite this, we found a *subset* of release sites was enriched with NMDARs, and modeling of glutamate release and receptor activation in measured synapses indicated this nanotopography promotes NMDAR activation. This subset of release sites was internally denser with Munc13-1, aligned with abundant PSD-95, and associated closely with specific NMDAR nanodomains. Further, NMDAR activation drove rapid reorganization of this release site/receptor relationship, suggesting a structural mechanism for tuning NMDAR-mediated synaptic transmission. This work reveals a new principle regulating NMDAR signaling and suggests that synaptic functional architecture depends on the assembly of and trans-cellular spatial relationships between multiprotein nanodomains.

## INTRODUCTION

Many cellular functions are performed by macromolecular protein ensembles that require nanoscale spatial relationships with neighboring ensembles to facilitate complex signaling.

These critical relationships are required in healthy signaling and disrupted in disease across biological systems. This is especially clear in the neuronal synapse^1–3^. Synapses mediate highly complex intercellular signaling to respond extremely rapidly and with high fidelity to diverse stimuli, propagating information in the brain. Despite their small size (<500 nm), the spatial position of signaling events *within* the synapse can critically influence their effect on neuronal signaling^4–7^. Indeed, converging lines of evidence suggest the nanoscale positioning, relative to one another, of the protein ensembles that mediate synaptic signaling is a key determinant of local synapse function^8^.

A clear case of how nanoscale coordination of protein ensembles influences synaptic transmission is the regulation of ionotropic glutamate receptor activation. Presynaptic vesicle release machinery, postsynaptic receptors, and the scaffolds that position them each concentrate in <100 nm diameter subsynaptic regions of high protein density (nanoclusters, NCs)^8^. Alignment of NCs across the synapse from one another into the trans-synaptic “nanocolumn” plays a critical role in regulating synaptic strength by enriching AMPA receptors (AMPARs) across the synaptic cleft from release sites^9,10^. Indeed, perturbing this complex nanoscale relationship between AMPARs and release sites disrupts synaptic transmission^1,11–13^, most remarkably even if synaptic receptor content is not altered^14^, demonstrating the fundamental role of nanoscale protein contextual relationships in neuronal function.

Despite the importance of nanoscale context for AMPAR function, how synaptic NMDA receptors (NMDARs) are organized relative to release sites remains unanswered. NMDARs are required for learning and memory and disrupted in many neurological disorders^15^, highlighting the need to understand their regulation. Importantly, though, the molecular context of NMDARs is critical for determining synaptic function even beyond controlling their activation, as the receptors form large signaling super-complexes that are intimately involved in both driving plasticity and establishing molecular organization within synapses^15–17^. Indeed, NMDARs signal via both Ca^2+^ flux-dependent and independent mechanisms^16,18^, and are attached via their large extracellular domains and long C-terminal tails to myriad extra and intracellular signaling proteins by which they are presumed to organize subsynaptic signaling domains^19–21^. These roles for NMDARs also depend on receptor subunit composition: receptor kinetics and biophysics are subunit specific^16^, super-resolution microscopy has shown differences in subunit distribution within single synapses^22,23^, and GluN2 subunit interactions with scaffolds and signaling molecules are differentially mediated by subunit C-tails^19,20,24^. It is therefore critical to understand NMDAR subunit positioning and molecular context, as these key attributes will control receptor activation and may also indicate regions where NMDARs create their own unique environments to facilitate subsynaptic signaling. However, we have until recently lacked the tools to investigate multiple protein complex relationships simultaneously, and therefore lack basic maps of receptor context to aid these determinations.

To map the spatial relationships of endogenous NMDAR GluN2 subunits to key pre and postsynaptic nanodomains in synapses, we utilized the multiplexing capabilities of DNA Exchange-PAINT (Points Accumulation for Imaging in Nanoscale Topography). We targeted the relationships in cultured rat hippocampal neurons between the critical GluN2 subunits GluN2A and GluN2B^15^, the major NMDAR postsynaptic scaffold protein PSD-95^25^, and presynaptic release sites, as marked by NCs of the vesicle priming protein Munc13-1^26–29^. This multiplexed mapping approach revealed nanodomain assembly principles in single synapses. Most surprisingly, we find that NMDARs are typically lacking from the immediate nanoscale region of the PSD directly across from presynaptic release sites. However, a subset of release sites was located near GluN2 subunits, and modeling based on the measured protein locations showed this topography promoted NMDAR activation. Consistent with multiple release site populations, the protein content and trans-cellular molecular context of individual presynaptic release sites in single synapses differed markedly from one another. Only a subset of Munc13-1 NCs were enriched across the synapse with high-density PSD-95, and these nanocolumnar NCs had significantly higher Munc13-1 protein density and were dramatically more enriched with GluN2 subunits than non-nanocolumnar release sites in the same synapses. Further, NMDAR activation increased their enrichment to nanocolumnar, but not non-nanocolumnar, release sites, suggesting that plasticity of nanoarchitecture may help regulate the subset of NMDARs activated during neurotransmission. These results reveal that NMDAR positioning and organization are governed by specific trans-cellular spatial relationships between multiprotein ensembles and suggest that overall synapse architecture may arise from local formation of subsynaptic domains with unique functional roles.

## RESULTS

### Mapping endogenous NMDAR organization with DNA-PAINT

We took advantage of the high resolution and multiplexing flexibility of the single molecule localization microscopy (SMLM) technique DNA Exchange-PAINT^30^ to map the nanoscale relationships of multiple proteins at a single synapse. To facilitate multiplexing, we preincubated^31,32^ primary antibodies separately with secondary nanobodies conjugated to orthogonal DNA-PAINT docking strands, then combined them on the sample (Fig. 1a, and see Materials and Methods). This method allows simple, antibody species-independent multiplexing and saturates the primary antibody with a defined number of DNA strands close to ideal for SMLM (1 per nanobody; 2 nanobodies per antibody). The use of small secondary nanobodies also significantly reduces linkage error compared to a secondary antibody^31,32^ and avoids the technical hurdles of direct primary antibody conjugation. NC-sized (40-100 nm) regions of high protein density can be observed within single synapses (∼300-500 nm) in dendritic spines (∼1 µm) in DNA Exchange-PAINT renderings of GluN2A, the pre-and post-synaptic scaffolds Bassoon and PSD-95, and a myristoylated-EGFP cell fill (Fig. 1b-d), revealing the power of this technique to resolve the nanoscale organization and context of multiple proteins at a single synapse.

**Figure 1.**
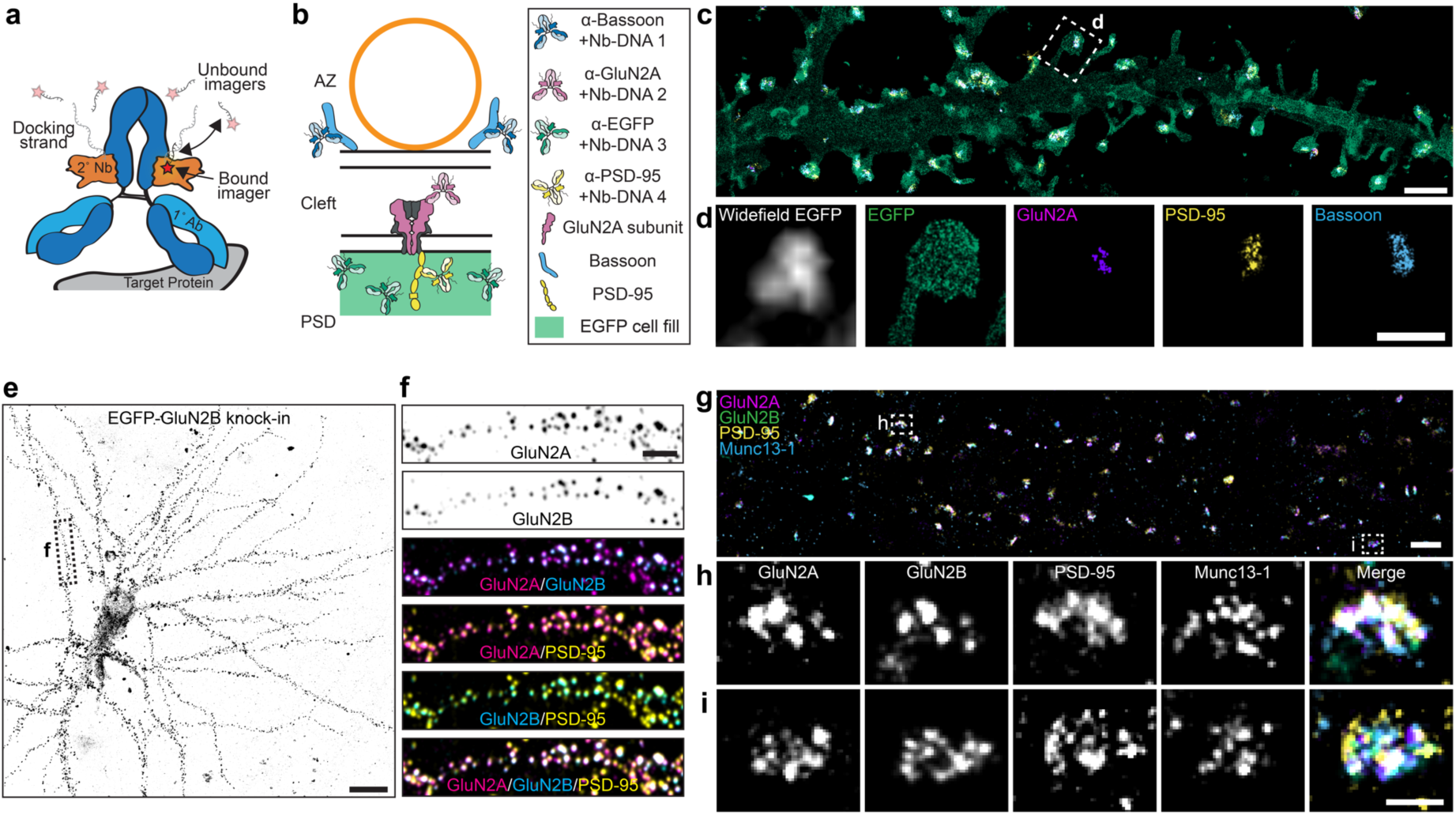
Mapping endogenous NMDA receptor organization with DNA-PAINT **a** Schematic of a primary antibody labeled with a DNA-PAINT docking strand-conjugated secondary nanobody and imaged with fluorescent imager strands. Red star indicates fluorophore. **b** Schematic of four-target DNA-PAINT labeling using primary antibodies preincubated with secondary nanobodies. **c** DNA-PAINT rendering (10 nm pixels) of myristoylated-EGFP cell fill, surface expressed GluN2A, PSD-95, and Bassoon demonstrating four-target, synaptic DNA-PAINT. Scale bar 2 µm. **d** Boxed region from top, including widefield image of myr-EGFP. Scale bar 500 nm. **e** Confocal image of EGFP-GluN2B CRISPR knock-in cell. Scale bar 20 µm. **f** Boxed region from left, showing surface expressed GluN2A, surface expressed GluN2B (EGFP knock-in), and PSD-95 labeling colocalized at synapses. Scale bar 4 µm. **g** DNA-PAINT rendering (10 nm pixels) of endogenous, surface expressed GluN2A and GluN2B (EGFP knock-in), PSD-95, and Munc13-1. Scale bar 1 µm. **h-i** Zoom-in on two representative synapses showing nanoclusters of each protein and their co-organization. Scale bar 200 nm.

We aimed to measure the distribution of endogenous, surface expressed NMDAR subunits with both pre-and postsynaptic molecular context. To do this, we performed 2D DNA Exchange-PAINT of GluN2A, GluN2B, PSD-95 and Munc13-1 together at synapses in DIV21 rat primary hippocampal cultures. We labeled surface-expressed GluN2A (Supplementary Fig. 1a-c) and total PSD-95 and Munc13-1 with antibodies to avoid overexpression artifacts. As we were unsatisfied with commercially available antibodies targeting surface expressed GluN2B, we used ORANGE CRISPR^33^ to knock in EGFP to the extracellular domain of endogenous GluN2B, which tolerates N-terminal tags well; in particular EGFP is known to have minimal effect on GluN2B function or distribution when inserted in similar locations^33–35^. We then labeled surface-expressed EGFP-GluN2B with an anti-EGFP antibody and visualized its synaptic distribution by confocal microscopy (Fig. 1e-f and Supplementary Fig. 1d). Surface-labeled EGFP-GluN2B was, as expected and consistent with previous work^33^, extensively colocalized at synapses with PSD-95, suggesting the knock-in receptor trafficked appropriately. We were routinely able to identify synapses containing all four proteins (Fig. 1g-i), and only analyzed synapses containing EGFP-GluN2B localizations, as those without EGFP-GluN2B may either genuinely lack the subunit or could have accumulated an indel during genome editing and be GluN2B knock-outs. We further only selected synapses perpendicular (en face) to the optical axis for further analysis to best measure positional relationships between proteins with high fidelity (Supplementary Fig. 2). Cross-talk between these docking strands and fluorophores is quantitatively extremely limited^36,37^, and lack of cross-talk was apparent in the very different distributions of, for example Munc13-1 and PSD-95 (Fig. 1g-i, Fig. 5A) or GFP cell fill and Bassoon (Fig. 1c-d), which were imaged simultaneously. Together, these data demonstrate a workflow to map the distribution of surface-expressed, endogenous NMDAR subunit nanoclustering and position in the spatial context of other key pre and postsynaptic nanodomains.

### Endogenous GluN2 subunits form diverse nanodomain types

In pyramidal neurons in hippocampus and neocortex, the dominantly expressed GluN2A and GluN2B subunits^15^ are each found in subsynaptic NCs that are postulated to control receptor subtype-specific activation and activity-regulated positioning^22,23,38^ relevant for signaling and plasticity. The characteristics and mutual relationships of these receptor-containing areas may shape their relationships with other synaptic constituents. Therefore, before investigating the spatial organization of NMDARs relative to other nanodomains, we first examined the nanoscale organization of GluN2A and GluN2B.

As expected^23^, both GluN2A and GluN2B formed small, tight NCs within the synapse that were readily discernible in local density heat maps (Fig. 2a; n.b. we use “NC” throughout to refer to subsynaptic nanoclusters, not to the nanocolumn (i.e. aligned pre/post NCs)).To characterize these NCs, we first measured the normalized autocorrelation^10^ of each subunit (Fig. 2b), which indicates the length scales over which the density of a protein correlates to itself, normalized to a uniform distribution at the average density. The autocorrelation thus assesses nanoclustering without needing to define NC boundaries, and an autocorrelation of one indicates the protein of interest is randomly distributed^10^. Both GluN2A and GluN2B showed autocorrelation magnitudes greater than one at short length scales that quickly decayed and plateaued after 35-45 nm, consistent with each protein forming NCs near that diameter. Both autocorrelations also plateaued below one, consistent with most of the protein being concentrated within NCs and sparse between, as was visually apparent (Fig. 2a). This can be contrasted with the broader autocorrelation for PSD-95, reflecting the known larger size of its NCs and stronger presence of PSD-95 nearly continuously between NCs^10,39–41^.

**Figure 2.**
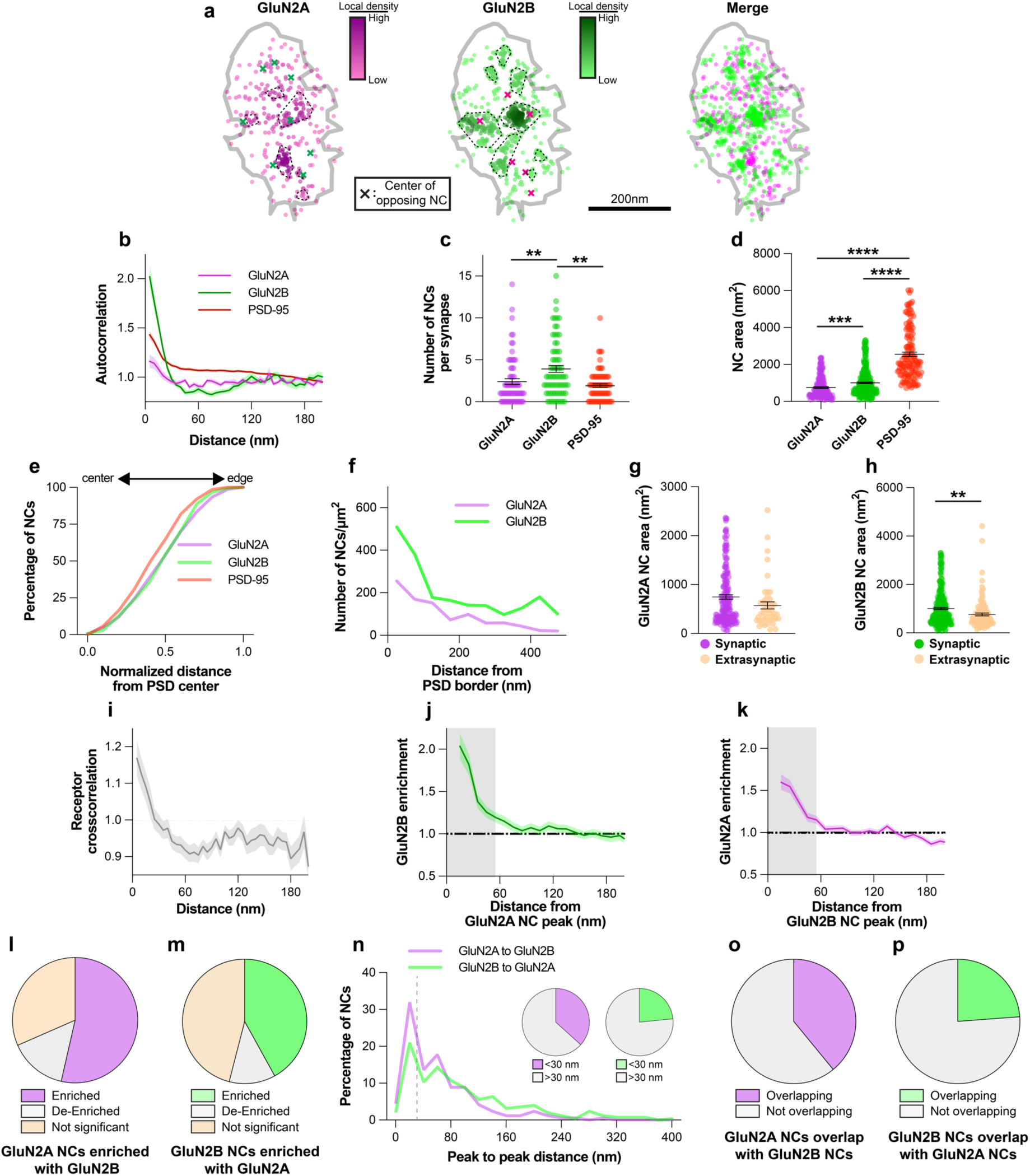
Endogenous GluN2 subunits form diverse nanodomain types **a** Example synapse of DNA-PAINT localizations of GluN2A and GluN2B showing GluN2 NCs have diverse co-organization. Each point is a localization, and its heat map codes normalized local density. NCs are indicated by dash-bordered areas. Peaks of the NCs of the opposing protein are indicated by colored x’s. Gray outline indicates PSD border, defined by PSD-95 localizations (not shown). **b** GluN2 subunit autocorrelations decayed faster than PSD-95 and plateaued below one, indicating small NCs with few localizations between them. **c** GluN2B NCs were more numerous than GluN2A or PSD-95 NCs. **d** Both GluN2A and GluN2B NCs were smaller than PSD-95 NCs. **e** GluN2 subunit NCs were distributed slightly less centrally than PSD-95 NCs. **f** Extrasynaptic GluN2 tended to be within ∼200 nm of the PSD edge. **g-h** Extrasynaptic GluN2 clusters were on average smaller than synaptic GluN2 NCs. **i** Cross-correlation and **j-k** cross-enrichments indicated strong overlap of GluN2A and GluN2B densities at short distances. **l-m** 52.6% of GluN2A and 42.0% of GluN2B NCs were significantly enriched with the opposite subunit. **n** 36.7% of GluN2A and 22.6% of GluN2B NC peaks were located within 30 nm of an opposite GluN2 NC peak**. o-p** 39.0% of GluN2A and 23.4% of GluN2B NC areas spatially overlapped with the opposite subunit. Data in **b** and **i-k** are means ± SEM shading. Points in **c** are individual synapses and points in **d** and **g-h** individual NCs. Lines in **c, d, g** and **h** are means ± SEM. Data in **n** are shown as frequency histograms (20 nm bins), with dashed line indicating the division summarized in inset pie charts. N = 173 GluN2A, 274 GluN2B, 134 PSD-95 NCs from 74 synapses throughout synaptic data; N = 49 GluN2A and 119 GluN2B extrasynaptic NCs from 74 synapses. Specific n per data point in **b** and **i** varies from 74 due to data processing, see methods for details. *p<0.05, **p<0.01, ****p<0.0001

We measured the properties of individual NCs by identifying them directly using DBSCAN (density-based spatial clustering of applications with noise)^42^. We identified on average 2.4 ± 0.3 GluN2A and 3.9 ± 0.4 GluN2B NCs per synapse, with average areas of 748 ± 44 nm^2^ and 1006 ± 45 nm^2^, respectively, consistent with the size of clustering suggested by the autocorrelation (∼30-40 nm diameters if approximating a circular NC) (Fig. 2c-d). These NCs were clearly different in size and number from the larger, less frequent PSD-95 NCs identified at the same synapses. We also observed, using 2D STED, that both subunits formed visually similar NCs with similar size to DNA-PAINT (Supplementary Fig. 3), recapitulating their basic organization with an independent super-resolution technique and labeling method. GluN2 NCs were distributed throughout the radial extent of the synapse, compared to PSD-95 NCs that skewed slightly more central (Fig. 2e). As has been reported also in tissue slices^43^, some synapses had centrally located NMDAR NCs, whereas others had prominent NCs positioned near the PSD edge. We also observed GluN2 NC-sized objects in extrasynaptic regions. These were largely concentrated within ∼200 nm of the edge of the synapse (Fig. 2f), consistent with recent single particle tracking data of expressed receptor subunits^44^, and on average were slightly smaller than their respective synaptic NCs (GluN2A: 574 ± 74 nm^2^, p=0.0673 vs synaptic; GluN2B: 767 ± 58 nm^2^, p=0.0025 vs synaptic; Fig. 2g-h).

NMDARs of different GluN2 subunit compositions may form synaptic nanodomains in which separate diheteromeric receptor types (diheteromers) intermix, or they could represent triheteromeric GluN1/2A/2B receptors (triheteromers), or be a combination of both. Given the unique kinetics and interactors of each subunit, these mixed subunit nanodomains could have unique activation properties or downstream signaling pathways depending on their constituents and proximity to other complexes. We therefore assessed the nanoscale relationship of endogenous GluN2 subunits to one another. We first measured the relationship of receptor subunits using a normalized cross-correlation^10^, which indicates the spatial scales over which two protein distributions correlate. Like the autocorrelation, the cross-correlation is normalized to the cross-correlation of two uniformly distributed proteins; a magnitude of one indicates the relationship is not different from random^10^. The magnitude of the cross-correlation of GluN2A and GluN2B was greater than one at length scales less than 30 nm (Fig. 2i), similar to the predicted size of GluN2 NCs (Fig. 2b,d), indicating that on average, GluN2A and GluN2B NCs are strongly spatially associated. Indeed, a cross-enrichment analysis^10^, which measures the density of one protein surrounding the peak density position of each NC (peak) of another protein (similarly normalized to a randomized distribution of one protein such that a magnitude of one is not different from random), showed that each subunit was on average significantly enriched near the opposite subunit’s NC peak (Fig. 2j-k). When the cross-enrichment curves were separated into those that statistically can be determined as enriched or de-enriched within the first 60 nm from the peak^10^, we found that 53.5% of GluN2A NCs were enriched with GluN2B and 41.9% of GluN2B NCs were enriched with GluN2A. By contrast, only 14.9% of GluN2A and 12.0% of GluN2B NCs were statistically *de*-enriched with the other subunit (Fig. 2l-m). Consistent with this finding, 36.7% of GluN2A NC peaks could be found within 30 nm of the nearest GluN2B NC peak, and 22.6% of GluN2B NC peaks were within 30 nm of the nearest GluN2A peak (Fig. 2n). Further, 39.1% and 23.8% of GluN2A and GluN2B NC areas, respectively, spatially overlapped (Fig. 2o-p). These data together indicate that 30-50% of GluN2A and GluN2B NCs are closely spatially associated with one another at these synapses, a number strikingly similar to the predicted percentage of triheteromeric NMDARs at mature synapses^45,46^.

Finally, we took advantage of STED to ask whether steric hindrance between the various probe molecules might limit our interpretations of the spatial relationship between subunits. We compared the distance between GluN2A and GluN2B NC peaks when GluN2B was labeled with probes of different size while GluN2A labeling was held constant; if sterics were limiting, this distance should be larger when using larger probes. As expected, GluN2A NCs were the same size between GluN2B labeling conditions (Supplementary Fig. 3g) and we observed smaller GluN2B NCs when using a smaller probe (Supplementary Fig. 3h). Critically, however, the nearest-neighbor distance between GluN2A and GluN2B NCs was not different between conditions (Supplementary Fig. 3i), suggesting our analyses were not significantly limited by sterically hindered access of probes to the target proteins. These results suggest subtype-specific NMDAR trafficking mechanisms establish a diverse array of nanodomain types within single synapses, where the co-enriched population likely represents either triheteromers or mixed nanodomains of GluN2A and GluN2B diheteromers, while the remaining NCs likely represent nanodomains that accumulate a single diheteromer type.

### Only a subpopulation of release sites is enriched with GluN2 subunits

NMDARs with different subunit compositions have different sensitivity to distance from the release site^47^, and their position within the synapse may affect what intracellular signaling cascades are activated after Ca^2+^ influx. To examine the relationship of GluN2 subunits to release sites, we detected Munc13-1 NCs, which have been hypothesized to physically mark vesicle docking and fusion sites within individual mammalian synapses^29^ and the drosophila neuromuscular junction^26,28^. The Munc13-1 autocorrelation sharply decayed and plateaued below one around 45 nm, indicating small NCs with few localizations between (Fig. 3a-b). This was supported by direct NC detection, which showed a wide range of NC numbers per synapse (up to 17, mean 6.8 ± 0.5 with area 851 ± 28 nm^2^, Fig. 3c), consistent with previous reports^29,48^. In contrast to our expectations, GluN2A and GluN2B were each quite strongly *de-*enriched from Munc13-1 NC peaks (Fig. 3d-e). Nearest the peak, GluN2A enrichment was only 0.76 ± 0.04, and GluN2B enrichment 0.58 ± 0.04, of randomly distributed receptor density, and their enrichments did not reach random until 95 and 155 nm from the Munc13-1 NC peak, respectively. Further, the GluN2 enrichment indices, or the average enrichment within 60 nm of the Munc13-1 NC peak^10^, was 0.87 ± 0.03 and 0.77 ± 0.03 for GluN2A and GluN2B, respectively, indicating a significant lack of both subunits immediately across the synapse from Munc13-1 NC peaks. This surprising observation provides direct evidence that NMDAR distribution within synapses is sensitive to presynaptic organization.

**Figure 3.**
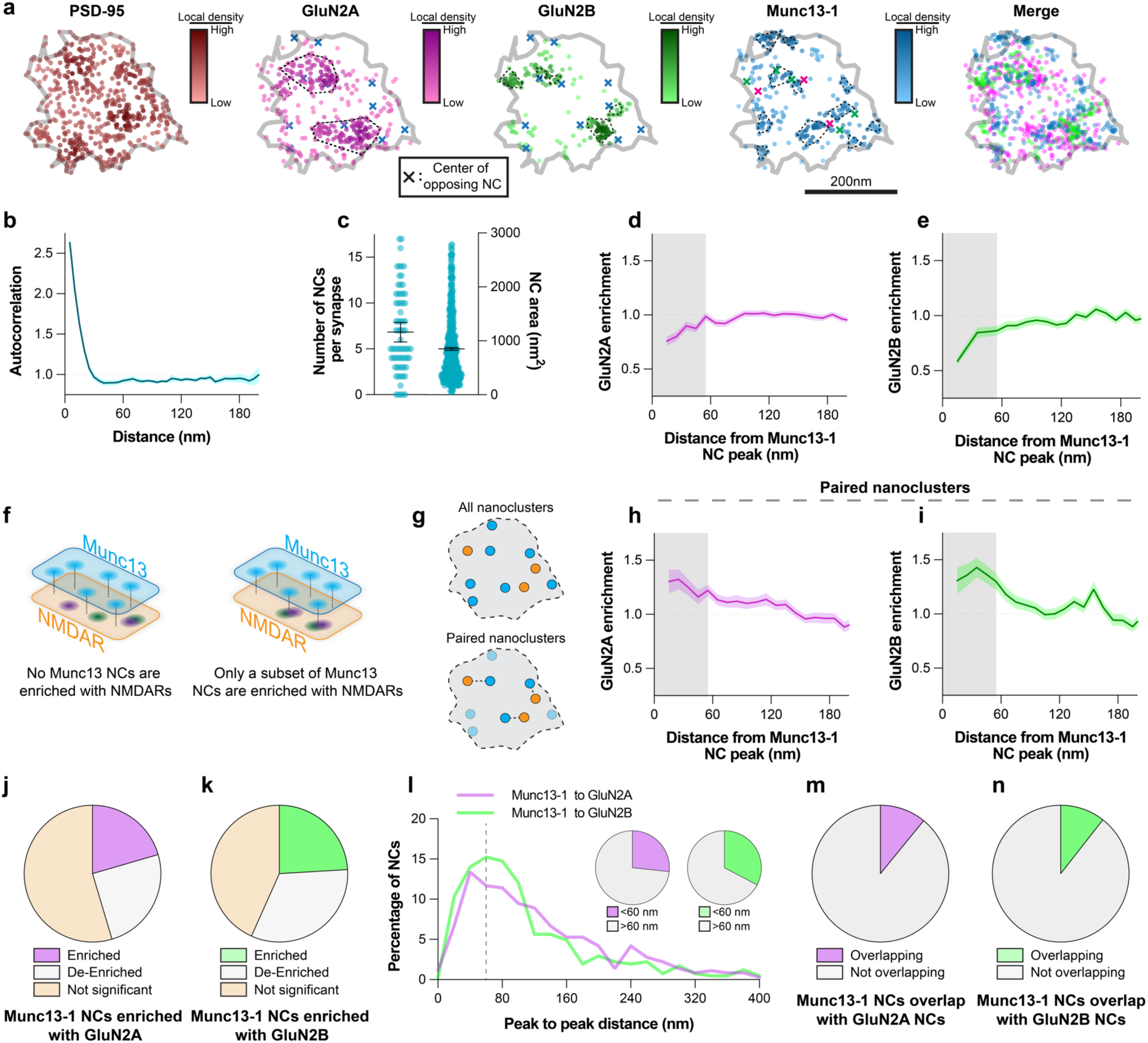
Only a subpopulation of release sites is enriched with GluN2 subunits **a** Example synapse of DNA-PAINT localizations of PSD-95, GluN2A, GluN2B, and Munc13-1 showing arrangement of receptor subunits relative to release sites. Markers as described in Fig. 2a. **b** Munc13-1 autocorrelation decayed rapidly and plateaued below one, consistent with its **c** numerous and small detected Munc13-1 NCs. **d-e** Munc13-1 NCs were, on average, de-enriched with GluN2A and GluN2B at distances <55 nm (shading). **f** Schematic indicates possible configurations of GluN2 and Munc13-1 NCs. **g** Schematic of NC pairing to reveal stereotyped distances of closely-associated NCs. **h** Paired Munc13-1 NCs were enriched with GluN2A as well as **i** GluN2B within 55 nm of their peak. **j-k** 20.5% and 24.1% of Munc13-1 NCs were statistically enriched with GluN2A or GluN2B. **l** 26.7% and 32.7% of Munc13-1 NCs had a GluN2A or GluN2B NC peak within 60 nm. **m-n** 10.9% and 10.6% of Munc13-1 NCs spatially overlapped with GluN2A or GluN2B NCs. Data in **b**, **d-e, and h-i** are means ± SEM shading. Points in **c** (left) are synapses and (right) NCs, with lines at mean ± SEM. Data in **l** are shown as frequency histograms (20 nm bins), with dashed line indicating the division summarized in inset pie charts. N = 173 GluN2A, 274 GluN2B, 478 Munc13-1 NCs from 74 synapses throughout, except **h-i**, where N = 113 Munc13-1 NCs when paired with GluN2A and 152 when paired with GluN2B. Specific n per data point in **b** varies from 74 due to data processing, see methods for details. *p<0.05, **p<0.01, ***p<0.001.

The offset from release sites was intriguing given that it would appear to decrease the likelihood of NMDAR activation, particularly of receptors containing GluN2B, which was on average more strongly de-enriched from Munc13-1 NCs and also has a stronger predicted distance-dependence for activation due to its slower glutamate binding^47^. However, having observed diversity in the spatial relationship between GluN2 subunits, we asked whether the relationship of GluN2 subunits to Munc13-1 NCs was a result of a systematic de-enrichment of receptors around all Munc13-1 NCs, or if there existed a subset of release sites enriched with GluN2 subunits (Fig. 3f). Such diversity would suggest a tight interplay of NMDAR position and active zone structure.

As a first step, we paired mutually nearest Munc13-1 and GluN2 NCs and measured their cross-enrichment (Fig. 3g). If GluN2 density were systematically de-enriched from Munc13-1 NCs, then it would remain de-enriched even after selecting for nearest pairs. However, we observed that paired Munc13-1 NCs were significantly cross-enriched with both subunits, consistent with there being a subpopulation of release sites enriched with NMDARs (Fig. 3h-i). Indeed, even without pairing NCs, 20.5% and 24.1% of Munc13-1 NCs were statistically enriched with GluN2A and GluN2B, respectively, despite a slightly larger population being statistically de-enriched (24.9% for GluN2A and 32.6% for GluN2B, Fig. 3j-k). Further, 26.7% and 32.7% of Munc13-1 NC peaks had a GluN2A or GluN2B NC peak, respectively, within 60 nm (Fig. 3l), and 10.9% and 10.6% of Munc13-1 NCs showed spatial overlap with GluN2A and GluN2B, respectively (Fig. 3m-n). Intriguingly, while 34.0% of the release sites enriched with GluN2 subunits were specifically enriched with both GluN2A and GluN2B, just 6.3% were enriched with only one subunit and *de*-enriched with the other, suggesting triheteromer or mixed diheteromer nanodomains may be specifically enriched closer to release sites.

### NMDAR positioning near a subset of release sites regulates their probability of activation

To determine whether the diversity of this organization impacted synaptic function, we simulated glutamate release and the resulting synaptic subunit-specific NMDAR open probability (P_o_) at a representative subset of our measured synapses (Supplementary Table 5) using MCell^49,50^. We took advantage of the multiplexed nature of our DNA-PAINT dataset to define the spatial constraints and subunit locations for modeling (Fig. 4a): we defined the borders of the synapse using the combined localizations of PSD-95, Munc13-1, and GluN2A/B, identified the possible release site positions as the peak density position of each Munc13-1 NC, and simplified receptor positions as GluN2 subunit localizations, since receptor density is expected to be proportional to localization density.

**Figure 4.**
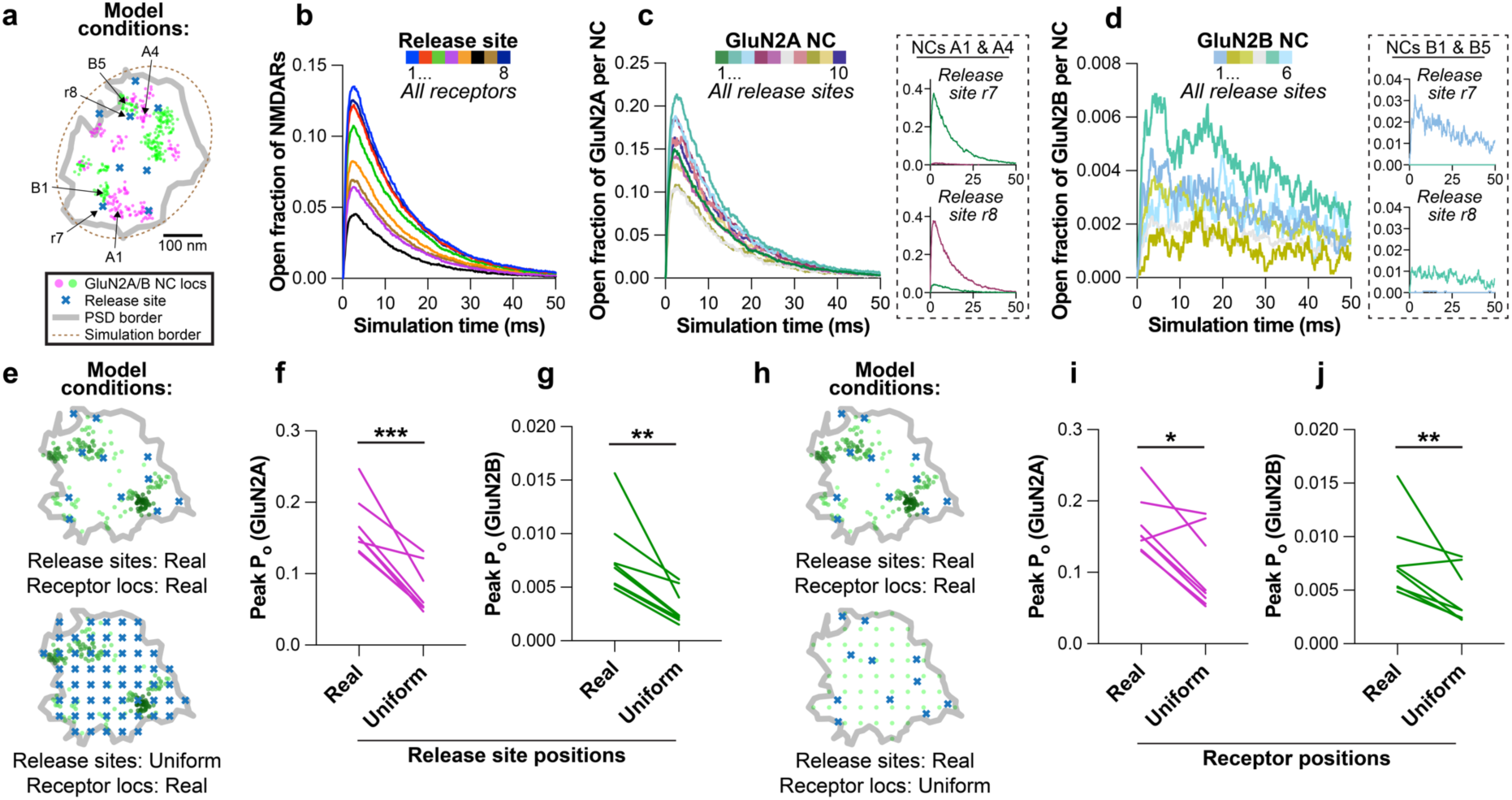
NMDAR positioning near a subset of release sites regulates their probability of activation **a** Schematic of receptor NC and release site positions in receptor activation modeling; synapse shown is exemplar used in **b-d**. Magenta and green circles indicate GluN2A and GluN2B localizations within NCs, respectively, blue x’s indicate release site positions, gray outline indicates PSD border, and dashed line represents minimal bounding ellipse used in modeling (see Methods). Location of release sites and NCs shown in **c-d** insets are indicated with labeled arrows. **b** The open fraction of all NMDARs at the synapse varies depending on where glutamate is released. **c-d** The open fraction of NMDARs varies by GluN2 NC for both GluN2A and GluN2B. Insets for each show two example NCs with differing P_o_ depending on which release site is active. **e** Schematic of receptor and release site positions compared in receptor activation modeling in **f-g**. Gray outline indicates PSD border, green circles indicate GluN2 localization positions, and blue x’s indicate release site positions. **f-g** Peak synaptic GluN2A and GluN2B open probabilities (P_o_) were significantly greater when release sites were in their real positions vs uniformly distributed. **h** Schematic of receptor and release site positions compared in receptor activation modeling in **i-j**. Symbols as in **e**. **i-j** Peak synaptic GluN2A and Glun2B P_o_ were significantly greater when receptors were in their real positions vs uniformly distributed. N = 8 synapses in **e-j** (data in **a-d** show one exemplar synapse from this dataset).

Examining single synapses, we found that the open fraction of total NMDARs (GluN2A + GluN2B) varied strongly depending on which release site was active (Fig. 4b), supporting a strong degree of transmission heterogeneity among release sites. Conversely, regardless of which release site was utilized, different fractions of both GluN2A and GluN2B were likely to open depending on the receptor NC they were in (Fig. 4c-d). Parsing this further, the distance-dependence of activation was particularly apparent in specific release site/GluN2 NC combinations – for example for GluN2A, release from site r7 caused robust activation of receptors in NC A1 but not NC A4, while release from site r8 strongly activated receptors in NC A4 but not NC A1; this was similarly apparent for GluN2B (Fig. 4c-d insets). This confirms the previously predicted subunit-specific sensitivity of activation^47^ here using measured receptor distributions.

To test the impact of nanostructure on NMDAR activation, we compared simulation results using the real protein distributions in 7 synapses to that from simulations when either the release sites or receptors were distributed within the synapse in a uniform pattern. Several outcomes were possible: 1) given the overall de-enrichment of release sites with receptors (Fig. 3d-e), we might expect the real distributions to yield *weaker* NMDAR P_o_ than a uniform distribution; 2) alternately, given the steep distance-dependence of activation especially of GluN2B^47^, the subset of release sites that are enriched or partially enriched with receptors (Fig. 3j-n) might be sufficient to yield a *stronger* response than a uniform distribution; 3) if the interaction of these biophysical and nanostructural features produces no net effect, then responses would not be altered by regularizing receptor or release positions. Our modeling results were consistent with the second scenario. NMDAR P_o_ was significantly higher for both GluN2A-and GluN2B-containing receptors when release sites were in their measured positions rather than uniformly distributed (Fig. 4e-g). Similarly, NMDAR P_o_ was higher for both receptor subunits when the receptors themselves were in their real positions rather than uniform (Fig. 4h-j). Together, these results indicate that despite their overall de-enrichment with one another, the relative spatial organization we observed between release sites and receptors likely impacts NMDAR activation and synaptic function by maximizing receptor opening around a subset of release sites.

### A subset of structurally unique Munc13-1 NCs is enriched with PSD-95 and in the nanocolumn

Our results suggest there may be unique, trans-cellular molecular contexts for some release sites that may influence receptor positioning. To resolve this, we examined the relative trans-synaptic enrichment of PSD-95 near Munc13-1 NCs (Fig. 5a). PSD-95 anchors receptors within the synapse^25^ and is a central component of the trans-synaptic nanocolumn^9,10,14^. However, there are approximately 3.5 times as many Munc13-1 NCs (Fig. 3c) as PSD-95 NCs (Fig. 2c), suggesting an architectural diversity that may be important for the control of receptor subsynaptic positioning. When all NCs were analyzed, Munc13-1 was weakly enriched on average across from PSD-95 NC peaks and PSD-95 was essentially randomly distributed across from Munc13-1 NCs, consistent with the large numerical mismatch (Fig. 5b). However, this average did not reflect a systematic or consistent offset between Munc13-1 and PSD-95, as identifying mutually paired NCs to even the numerical imbalance revealed strong, bidirectional enrichment (Fig. 5c).

**Figure 5.**
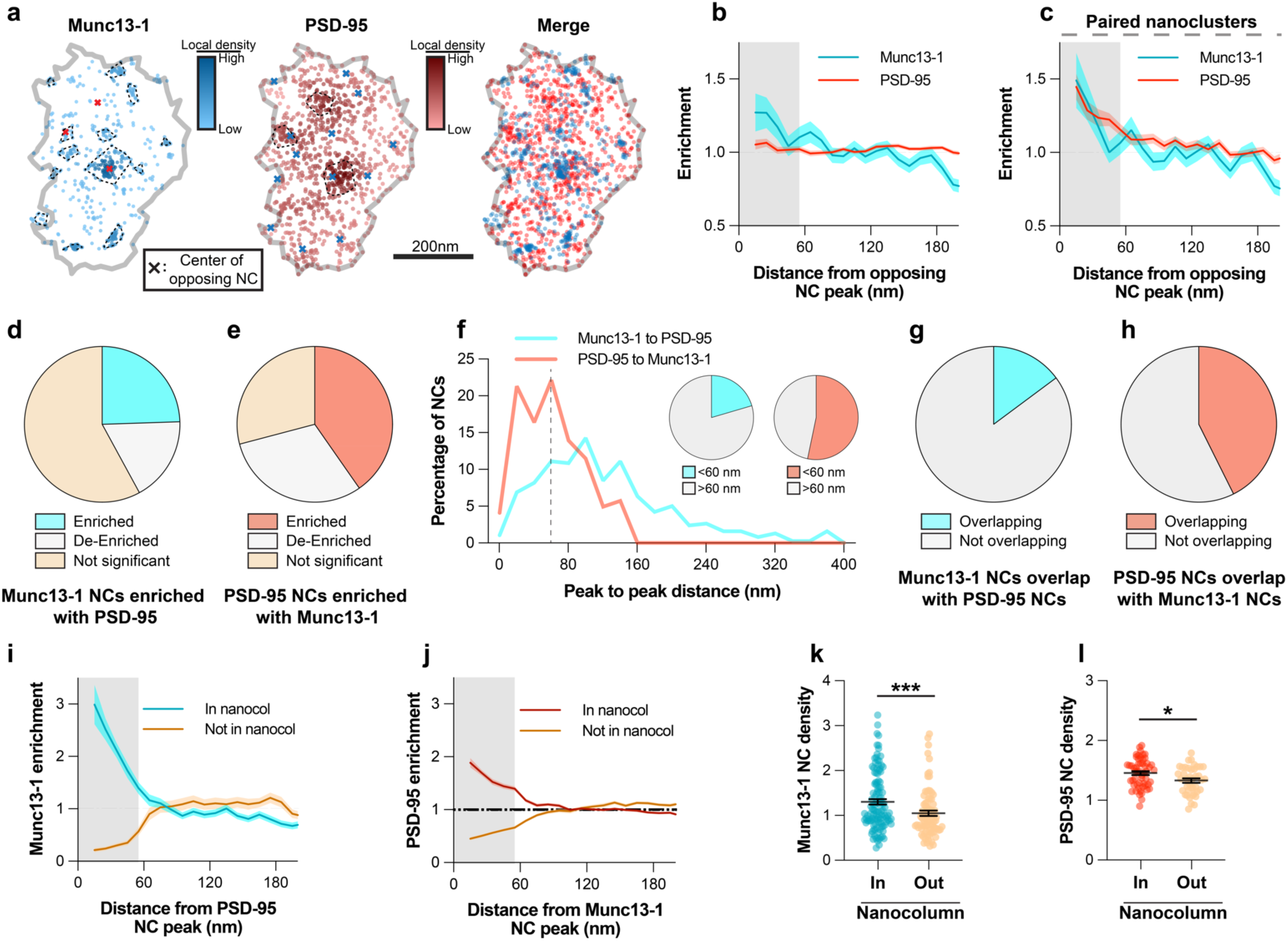
A subset of structurally unique Munc13-1 NCs is enriched with PSD-95 and in the nanocolumn **a** Example synapse of DNA-PAINT localizations of Munc13-1 and PSD-95, showing variable position of Munc13-1 NCs relative to PSD-95 density. Markers as in Fig. 2a. **b** On average, Munc13-1 and PSD-95 were weakly enriched with one another. **c** After pairing (as in **3g**), both PSD-95 and Munc13-1 were significantly enriched with one another. **d-e** 24.4% of Munc13-1 and 40.3% of PSD-95 NCs were statistically enriched with the other protein. **f** 20.5% of Munc13-1 and 53.3% of PSD-95 NCs had a NC peak of the other protein within 60 nm. **g-h**14.8% of Munc13-1 and 42.6% of PSD-95 NCs were spatially overlapped. **i-j** Cross-enrichments of PSD-95 NCs with Munc13-1 or vice versa demonstrate strong cross-enrichment of these proteins when subset by the data in **5d-e**. **k-l** Munc13-1 and PSD-95 NCs were denser when in the nanocolumn than when outside it. Data in **b-c** and **i-j** are means ± SEM shading. Points in **k-l** are NCs with lines at mean ± SEM. N = 174 PSD-95, 478 Munc13-1 NCs from 74 synapses in **b, d-h**. N = 98 paired PSD-95 and Munc13-1 NCs in **c**. N = 117 in nanocol, 84 not in nanocol Munc13-1 NCs; 54 in nanocol, 41 not in nanocol PSD-95 NCs from 74 synapses in **k-l**. ***p<0.0001, *p<0.05.

This subpopulation of Munc13-1 NC sites closely associated with PSD-95 was also evident in other measures. When we tested each NC for whether it was enriched with the other protein (within 60 nm), we found 24.4% of Munc13-1 NCs (∼1.5-2 per synapse, on average) were enriched with PSD-95 and 40.3% of PSD-95 NCs (∼1 per synapse, on average) were enriched with Munc13-1 (Fig. 5d-e). Additionally, 20.5% of Munc13-1 NCs and 53.3% of PSD-95 NCs had a nearest NC peak of the opposite protein within 60 nm (Fig. 5f), indicating a subpopulation of closely associated NCs. This fraction was similar to the proportion of each NC that spatially overlaid one another (14.8% for Munc13-1 NCs and 42.6% for PSD-95 NCs, Fig. 5g-h). These results together suggest that some Munc13-1 NCs have a privileged location closely associated with PSD-95 across the synapse. Indeed, this can be clearly seen in their cross-enrichment profiles (Fig. 5i-j) when subset by the data in Fig. 5d-e. We designate these mutually co-enriched PSD-95 and Munc13-1 densities as within the nanocolumn (nanocolumnar), which allowed us to make further conditional comparisons based on subsetting the data by nanocolumn status. Intriguingly, Munc13-1 NCs in the nanocolumn had a higher Munc13-1 density within 60 nm of their peak than those outside the nanocolumn (23.8% higher; 1.30 ± 0.06 inside vs 1.05 ± 0.06 outside, p=0.0003; Fig. 5k), and PSD-95 peak NC density was also significantly higher in the nanocolumn, though not to the same magnitude as Munc13-1 (9.0% higher; 1.46 ± 0.03 inside vs 1.33 ± 0.04 outside, p=0.0115; Fig. 5i). Together, these results suggest the nanocolumn represents a complex, macromolecular context that endows a privileged subset of release sites with high Munc13-1 NC density to spatially associate their neurotransmitter release most closely with PSD-95 NCs.

### Subunit-specific NMDAR nanodomains are organized with distinct trans-synaptic molecular contexts

With this approach in hand to molecularly classify subsets of Munc13-1 NCs, we hypothesized that nanocolumn organization may determine NMDAR positioning with respect to release sites. To test this, we made further conditional comparisons, assigning each Munc13-1 NC as being within the nanocolumn (enriched with PSD-95) or outside it (de-enriched with PSD-95), then measured GluN2 enrichment to these subsets. GluN2A was strongly de-enriched from Munc13-1 NCs outside the nanocolumn, considerably more so than to Munc13-1 NCs overall (enrichment index (EI): 0.60 ± 0.06 to Munc13-1 NC outside the nanocolumn vs 0.87 ± 0.03 to all Munc13-1 NCs (Fig. 3d); p=0.0012. However, GluN2A enrichment near Munc13-1 NCs *within* the nanocolumn was entirely rescued from de-enrichment, reaching a maximum enrichment of 1.23 at 55 nm (EI: 1.10 ± 0.68 to Munc13-1 NCs inside the nanocolumn vs 0.60 ± 0.06 to Munc13-1 NCs outside the nanocolumn, p<0.0001) (Fig. 6a-b, Supplementary Fig. 4a). Similar to GluN2A, GluN2B was strongly de-enriched from Munc13-1 NCs outside the nanocolumn (EI: GluN2B: 0.55 ± 0.06 vs GluN2A: 0.60 ± 0.06 to Munc13-1 NCs outside the nanocolumn, p=0.5922), but like GluN2A was significantly more enriched to Munc13-1 NCs in the nanocolumn (EI: 0.94 ± 0.08 to Munc13-1 NCs inside the nanocolumn vs 0.55 ± 0.06 to Munc13-1 NCs outside the nanocolumn, p=0.0069) (Fig. 6c, Supplementary Fig. 4b). These data indicate that while NMDARs are generally positioned away from release sites, they are significantly more associated with release sites in the nanocolumn.

**Figure 6.**
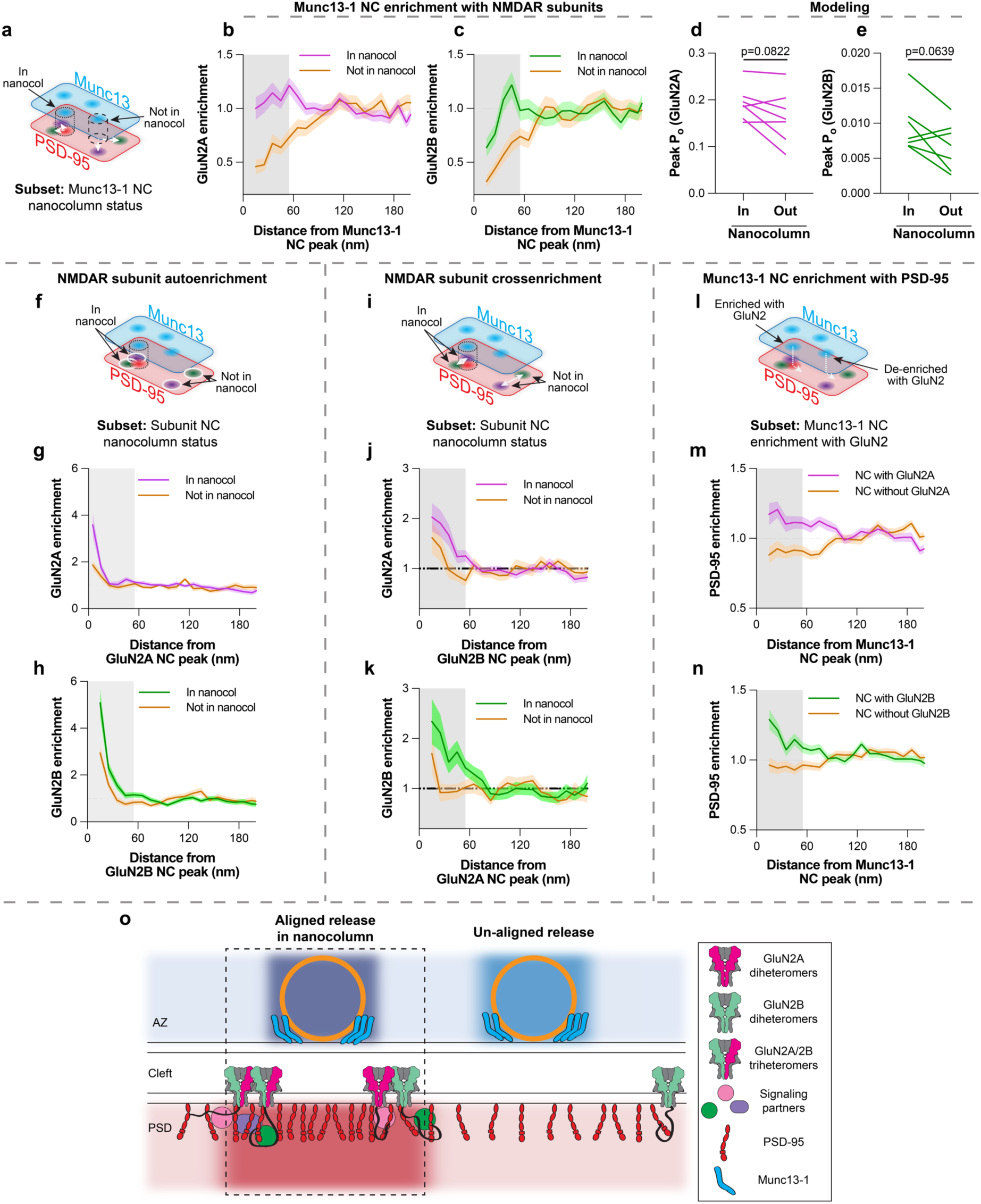
Subunit-specific NMDAR nanodomains are organized with distinct trans-synaptic molecular contexts In each section separated by dashed lines, the schematic (**a, f, i, l)** indicates the conditional comparison being made, and cross-enrichment plots (means ± SEM shading) show the measurements made with respect to GluN2A (**b, g, j, m**) and GluN2B (**c, h, k, n**). **a-c** Munc13-1 NCs in the nanocolumn were significantly more enriched with both GluN2A and GluN2B than Munc13-1 NCs outside the nanocolumn. N = 117 in nanocol, 84 not in nanocol Munc13-1 NCs from 74 synapses. **d-e** Receptor activation modeling showed glutamate release from nanocolumnar release sites trended toward greater synaptic activation of both GluN2A and GluN2B-containing NMDARs vs release from non-nanocolumnar sites. N = 7 synapses. **f-h** GluN2A and GluN2B NCs in the nanocolumn were denser than those outside the nanocolumn. N = 27 in nanocol, 21 not in nanocol GluN2A NCs; 42 in nanocol, 26 not in nanocol GluN2B NCs from 74 synapses. **i-k** GluN2A and GluN2B NCs were more cross-enriched with one another when in the nanocolumn. N as in **g-h**. **l-n** Munc13-1 NC enrichment with GluN2A and GluN2B can predict Munc13-1 NC enrichment with PSD-95. N = 96 Munc13-1 NCs with GluN2A, 119 without GluN2A, 115 with GluN2B, 156 without GluN2B NCs from 74 synapses. **o)** Model: NMDAR distribution in synapses is governed by nanodomains with distinct trans-synaptic molecular contexts. Active zones contain molecularly diverse release sites likely with differing vesicle priming and release properties and whose functional impact depends in part on differential trans-synaptic alignment. Receptor nanodomains near nanocolumn release sites contain GluN2A and GluN2B subunits; NMDARs accumulated near release sites will be activated more efficiently and potentially transduce unique signaling due to the presence of a mixed population of intracellular C-termini.

To untangle how these complex, conditional spatial relationships between proteins across the synapse from one another might impact NMDAR function, we returned to glutamate release modeling. Our measured organization and the modeling results in Fig. 4 suggest that release from nanocolumnar release sites should result in higher NMDAR P_o_. To test this prediction, we compared NMDAR P_o_ conditional on release from nanocolumnar release sites vs non-nanocolumnar release sites at the same subset of synapses as Fig. 4. Consistent with our prediction, the average NMDAR P_o_ trended higher when glutamate was released in the nanocolumn vs outside the nanocolumn (Fig. 6d-e) (GluN2A: p=0.0822; GluN2B: p=0.0639).

This difference was more pronounced for GluN2B-containing receptors, as expected for its lower glutamate affinity and steeper distance-dependence of activation^47^. These results suggest the conditional trans-synaptic spatial context of GluN2 subunits and release sites impacts NMDAR activation.

To pursue the molecular identity of these locations more fully, we next analyzed the characteristics of receptor NCs depending on their molecular context. Interestingly, both GluN2A and GluN2B NCs within the nanocolumn were denser compared to those outside the nanocolumn (Fig. 6f-h, Supplementary Fig. 4c-d), suggesting these receptor nanodomains may contain more receptors, or that the receptors are more tightly clustered near the nanocolumn. Because the enhanced accumulation was apparent for both subunits, we considered the possibility that the nanocolumn comprises a subdomain of specific receptor nanodomain types. We examined the cross-enrichments of each subunit to the other and found that GluN2A enrichment around GluN2B NCs was nearly 29% higher in the nanocolumn than out (Fig. 6i-j, Supplementary Fig. 4e; EI: 1.62 ± 0.16 inside nanocolumn vs 1.13 ± 0.17 outside of nanocolumn, p=0.0522). Even more strikingly, GluN2B enrichment around GluN2A NCs was enhanced almost 38% (Fig. 6k, Supplementary Fig. 4f; EI: 1.83 ± 0.23 inside nanocolumn vs 1.11 ± 0.15 outside of nanocolumn, p=0.0166). These observations suggest more abundant closely positioned subunits of each type within the nanocolumn. In further support of this, *none* of the nanocolumnar Munc13-1 NCs that were statistically enriched with any subunit were also *de-*enriched with the other. Instead, 37.3% of the nanocolumnar Munc13-1 NCs enriched with any subunit were significantly enriched with both subunits, with the remainder significantly enriched with one subunit and at least neutral with the other. This reveals a preferential positioning of enlarged, heterogeneous NMDAR nanodomains near release sites aligned with PSD-95, and more generally indicates that receptor subsynaptic organizational characteristics are dependent on trans-synaptic context.

Finally, we found the parallelized nanoimaging approach with DNA-PAINT further revealed that measuring the multiprotein context allows deeper and more accurate prediction of the organizational determinants of critical molecules. Notably, the presence of GluN2 density near a Munc13-1 NC was predictive of the postsynaptic environment around the release site: Munc13-1 NCs that were enriched with either GluN2A or GluN2B were on average also enriched with PSD-95, while those de-enriched with GluN2A or GluN2B were also de-enriched with PSD-95 (Fig. 6l-n, Supplementary Fig. 4g-h; EI of Munc13-1 NCs with PSD-95: 1.14 ± 0.04 with GluN2A vs 0.91 ± 0.03 without GluN2A, p<0.0001; 1.16 ± 0.03 with GluN2B vs 0.95 ± 0.03 without GluN2B, p<0.0001). These results together are consistent with a model of trans-synaptic nano-organization where GluN2 subunits are preferentially enriched near to, but offset from, nanocolumnar release sites to facilitate their activation (Fig 6o).

### Acute NMDAR activation drives rapid reorganization of release site/receptor relationship

If the nanoscale trans-synaptic protein context of NMDARs is a strong determinant of their activation, then NMDAR position relative to release sites is likely to be regulated by activity. To test this prediction, we asked whether the spatial organization of NMDARs and release sites was sensitive to activation of NMDARs themselves. We acutely (3 minutes) treated neurons with NMDA, a stimulation that strongly activates NMDARs^10,51^ and is capable of inducing nanoscale rearrangement of other synaptic proteins^10^, and focused on the relationship of GluN2A with release sites as it had a larger enrichment to nanocolumnar release sites (Fig. 6b).

While the concentration and duration of treatment we selected has previously been shown to not be excitotoxic^10,51^, we nevertheless first confirmed that NMDA treatment does not abrogate key nanostructure needed to evaluate spatial relationships after NMDAR activation. Nanoclustering of GluN2A, Munc13-1, and PSD-95 (Fig. 7a-f) was unchanged by NMDA treatment, as indicated by the similar autocorrelations between NMDA and vehicle (Fig. 7g-i), and while the number of Munc13-1 NCs was slightly higher in synapses treated with NMDA (Fig. 7j; 6.77 ± 0.40 vehicle, 8.48 ± 0.52 NMDA Munc13-1 NCs; p=0.0229), the distribution of release sites between nanocolumnar and not was unchanged (Fig. 7k-m). These results indicate that the core reference nanoscale relationships remain intact after NMDAR activation.

**Figure 7.**
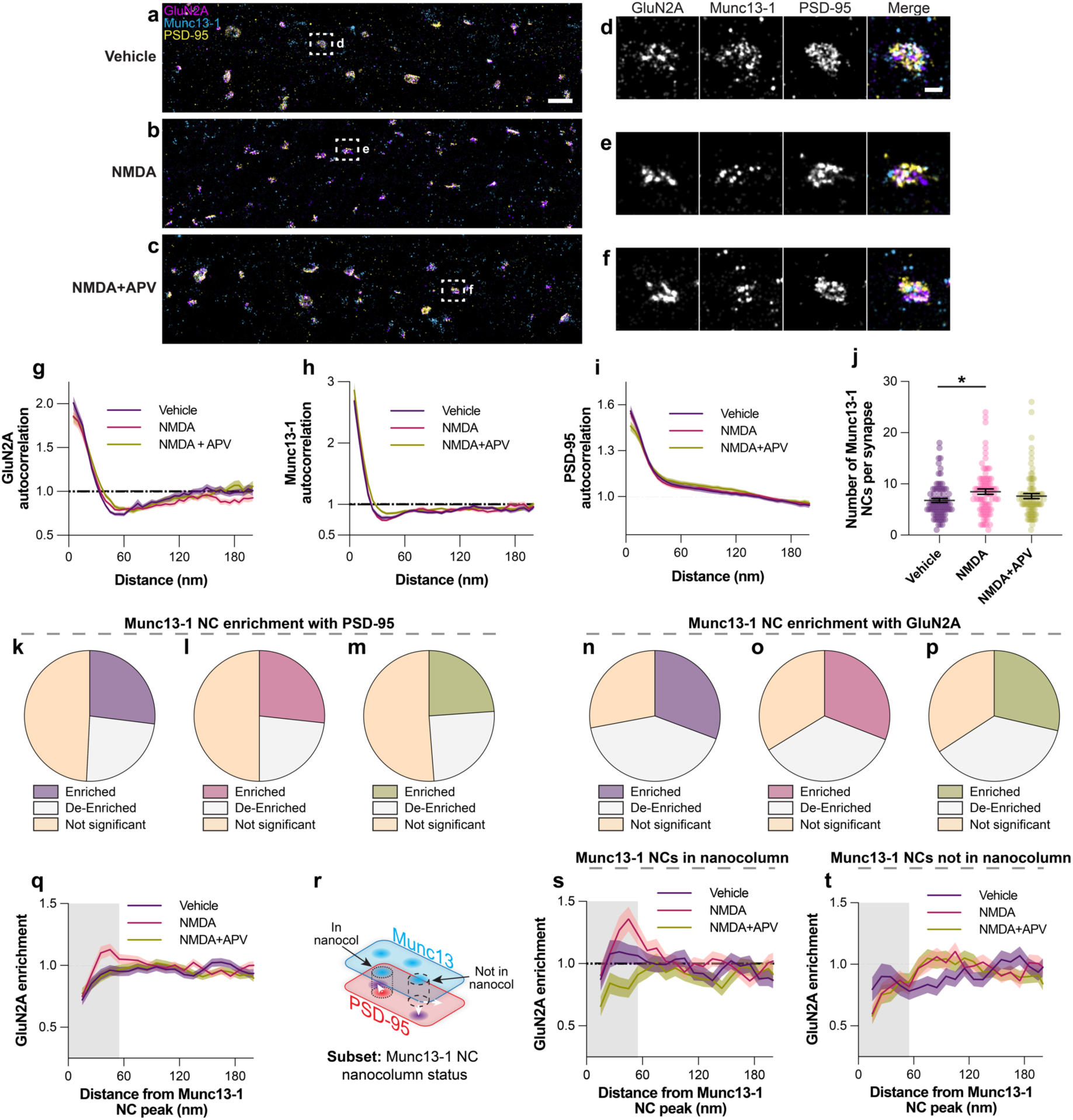
Acute NMDAR activation drives rapid reorganization of release site/receptor relationship **a-c** DNA-PAINT renderings of GluN2A, Munc13-1, and PSD-95 at synapses in neurons treated with vehicle, NMDA, or NMDA+APV. Scale bar 2 µm. **d-f** Zoom-in of boxed synapses in **a-c**. Scale bar 200 nm. **g-i** Autocorrelations of GluN2A, Munc13-1, and PSD-95 are not significantly altered by NMDA or NMDA+APV treatment, relative to vehicle. **j** The number of Munc13-1 NCs per synapse is slightly increased by NMDA treatment. N = 76 vehicle-treated, 80 NMDA-treated, and 82 NMDA+APV-treated synapses in **g-j. k-m** Subsetting of the cross-enrichment of Munc13-1 NCs with PSD-95 shows the percent of Munc13-1 NCs enriched with PSD-95 (i.e., in the nanocolumn) is not changed by NMDA or NMDA+APV treatment. **n-p** The percent of Munc13-1 NCs significantly enriched with GluN2A is not changed by NMDA or NMDA+APV treatment. **q** NMDA treatment selectively increases the average cross-enrichment of Munc13-1 NCs with GluN2A vs vehicle. This effect is rescued by co-application of APV. N = 508 vehicle-treated, 667 NMDA-treated, and 611 NMDA+APV-treated Munc13-1 NCs in **n-q**. **r** Schematic showing subsetting of Munc13-1 NC cross-enrichments with GluN2A by whether they are in the nancolumn or not. **s-t** NMDA treatment selectively increases the enrichment of Munc13-1 NCs with GluN2A within, but not outside, the nanocolumn. N = 137 vehicle-treated, 179 NMDA-treated, and 146 NMDA+APV treated nanocolumnar Munc13-1 NCs in **s**, and N = 121 vehicle-treated, 156 NMDA-treated, and 152 NMDA+APV-treated non-nanocolumnar Munc13-1 NCs in **t**. *p<0.05.

Notably, to avoid measuring any GluN2A that may have been internalized by NMDA treatment, we surface-labeled GluN2A after fixation (Supplementary Fig. 5) in this experiment rather than live as for previous experiments. This provided a serendipitous control for any influence of live antibody incubation on receptor clustering or internalization; the extremely similar autocorrelations, Munc13-1 NC number, and nanocolumn distribution of release sites between live (Fig. 2b, 3b-c, 5d) and fixed surface labeling (Fig. 7g-m) strongly indicate that live labeling did not significantly drive clustering or internalization of receptor subunits and lend confidence to the robustness of our measurements.

Finally, we evaluated the enrichment of Munc13-1 NC subsets with GluN2A. While the percentage of Munc13-1 NCs statistically enriched with GluN2A was not changed by NMDA (Fig. 7n-p), the average enrichment of a Munc13-1 NC with GluN2A was dramatically increased by acute NMDA treatment (Fig. 7q), peaking above 1, indicating that NMDAR activity was sufficient to drive rapid reorganization of this spatial relationship. This increase was specific to activation of NMDARs, as it could be prevented by co-application of the NMDAR antagonist APV, and NMDA+APV treatment did not itself affect protein clustering or the distribution of release sites into nanocolumns (Fig. 7a-m). Further, Munc13-1 NCs in vehicle-treated synapses were on average de-enriched with GluN2A (Fig. 7q), replicating this key distributional measure with the alternative fixation and staining method (compare vehicle to Fig. 3d). The greater average Munc13-1 NC enrichment with GluN2A after NMDA treatment appeared driven largely by nanocolumnar Munc13-1 NCs, which were more enriched with GluN2A after NMDA treatment, whereas the enrichment of non-nanocolumnar Munc13-1 NCs was unchanged (Fig. 7r-t). Given the lack of change in the percent of nanocolumnar and GluN2A-enriched Munc13-1 NCs, these data together suggest that NMDAR activation drives further GluN2A accumulation at existing nanocolumnar release sites already enriched with GluN2A and indicate that the trans-synaptic spatial organization of release sites and GluN2A is sensitive to NMDAR activation at a rapid timescale.

## DISCUSSION

We leveraged the high resolution and multiplexing capabilities of DNA Exchange-PAINT to map the nano-organization of GluN2A and GluN2B with respect to the key release site and scaffold proteins Munc13-1 and PSD-95. GluN2 NCs were only well aligned to release sites enriched with PSD-95 i.e. located at the nanocolumn, but otherwise were de-enriched from release sites. Biophysical simulations using super-resolved synaptic maps showed this organization promoted net receptor activation. This multi-protein relationship was plastic on a rapid time scale, suggesting a structural mechanism for tuning NMDAR-mediated synaptic transmission. Further, Munc13-1 NCs outnumbered PSD-95 and receptor NCs, varied substantially in their interior density of Munc13-1, and were in some cases quite far (100s of nm) from the nearest NMDAR NC, and our modeling showed variability in NMDAR responses within a single synapse depending on which release site was activated. This spatial segregation raises the possibility that independent fusion of vesicles at different release sites is likely to activate unique ratios of GluN2A or GluN2B-containing receptors, expanding the computational potential of the synapse.

A major question arising from these results is whether the structurally definable subsets of Munc13-1 NCs we have observed have functionally distinct release properties. One possibility is they have different preferred release modes (synchronous vs asynchronous and spontaneous vs evoked), which are suggested to be spatially segregated at the presynapse^52–56^. Release mode diversity is critically associated with NMDAR function, as functionally distinct pools of NMDARs respond more to spontaneous vs evoked release^57,58^ and activate unique downstream signaling cascades^59,60^. NMDARs may also be preferentially activated by asynchronous release^61^, though this has not been tested experimentally. As the molecular determinants establishing release site preference for one or more different release modes remain to be determined, we cannot yet determine whether any subsets we identified are more likely to participate in one or another mode. Our data also support the possibility that structurally distinct Munc13-1 NCs could quantitatively tune action potential-evoked release. The number of Munc13-1 NCs correlates strongly with the number of evoked release sites^29,62^, and our data along with recent results indicate that release sites within a single synapse have diverse molecular properties^6,62–64^. We found that Munc13-1 density was higher at nanocolumnar NCs than at those outside the nanocolumn. While the exact relationship between Munc13-1 content and release properties is not well understood, Munc13-1 density has been positively correlated with vesicle priming and release probability^62,65^. It is therefore tempting to speculate that Munc13-1 NCs at the nanocolumn may support higher P_r_ there than at other synaptic sites, possibly to further regulate frequency-dependent engagement during paired action potentials or sustained activity.

Our modeling results indicated that the relative organization of release sites and receptors we measured supports stronger subunit-specific activation than uniform organization. While some modeling based on the high affinity of NMDARs suggests a minimal effect of nanodomain organization on NMDAR activation^66^, taking the subunit-specific kinetic behavior into consideration suggests NMDAR activation is sensitive to release site position, especially for receptors containing GluN2B^47^. Because so few NMDARs are activated during typical responses^67^, this organization could help promote the relatively dependable activation of NMDARs during basal release by allowing nanocolumnar release sites to efficiently activate at least the small population of NMDARs nearby. Consistent with this, our modeling suggested nanocolumnar release was more effective at activating NMDARs than non-nanocolumnar release. Further, as discussed above, if denser Munc13-1 organization within nanocolumn release sites is indeed associated with higher release probability, this would likely accentuate the trends apparent in the modeling. Together these observations support the general idea that synaptic nanoarchitecture helps tune the strength and nature of NMDAR-mediated signaling.

We observed further that the receptor relationship to release sites is itself sensitive to activation of NMDARs, raising the possibility that neuronal control of this organization might regulate plasticity induction. Future experiments taking advantage of multiplexed super-resolution maps as a basis for functional modeling, even beyond numerical or biologically realistic but simulated positions^10,14,47,61,66^, may help to dissect the contribution of this mechanism in circuits.

Even when in the nanocolumn, the position of GluN2 peak enrichment was still offset relative to the presynaptic NC peak (55 and 45 nm for GluN2A and GluN2B respectively, Fig. 6). This is well beyond the expected linkage error in our primary antibody/secondary nanobody system^31,32^, and while steric constraints surely limit the ability of even preincubated antibodies to bind very closely spaced receptors in dense molecular environments, they did not appear to significantly limit our spatial interpretations. The observed offset is small enough that those nearest receptors will be activated at close to their maximum probability (which is still predicted to be fairly low, especially for GluN2B), but this distinctive organization may carry several other functions. Indeed, NMDARs act as synaptic signaling hubs^16,68^ that scaffold diverse downstream signaling molecules through their long C-terminal tails^20^. It’s possible that NMDAR maximal activation is balanced to allow for many or large binding partners to fit near scaffold nanodomains that concentrate further downstream signaling proteins. For example, the holoenzyme of the major GluN2 intracellular binding partner CaMKII is ∼27 nm in diameter^69^, a size that could be disruptive to fit into the dense PSD-95 NC environment. Another potential reason for this shift could be to allow space for AMPARs to access maximal activation, as suggested by Hruska et al.^22^, which is important not only because of their biophysical properties^70^, but also because their C-tails and auxiliary proteins, such as the PSD-95-anchoring stargazin/TARPγ2, are targets for CaMKII phosphorylation^71^. This model appears consistent with biochemical experiments showing that while TARPγ2, PSD-95, GluN2B C-tail, and CaMKII form phase condensates together, the highest concentrations of TARPγ2 and PSD-95 are spatially separated from those of GluN2B C-tail and CaMKII^72^. In either case, this offset organization seems likely to be facilitated by the large size of GluN2 C-tails (∼660 amino acids), which could result in the receptor channel and the extracellular domains mapped here being localized laterally quite far from their PSD-95-anchoring C-terminus^25^. While the structures of GluN2 C-tails remain unsolved and are presumably flexible in neurons^73^, high-resolution, multiplexed mapping of receptor intracellular and extracellular domains with their interacting proteins will provide additional insight to this organization. Note that it will also be important to explore whether the remaining population of NMDAR subunits that do not align with Munc13-1 NCs instead are anchored at other specific synaptic subdomains to maintain distinct functions or organize proteins there, or perhaps instead represent a mobile pool of diffusing receptors.

Further analysis of how NMDAR positioning is conditional on additional proteins should prove helpful in establishing the mechanisms that determine their distribution and role in the synapse.

Synaptic cell adhesion molecules likely also play a key role in organizing NMDARs^74^. Though the molecular players are better understood for AMPARs^1,9,14^, neurexins are likely involved, as specific splice variants of presynaptic neurexin-1 control NMDAR responses through interactions with cerebellins in hippocampus^75^. NMDARs themselves also have large extracellular domains stretching nearly across the synaptic cleft that can interact directly with adhesion molecules, such as neuroligin-1^76^ or EphB2^77^, and deletion of NMDARs is sufficient to rearrange Munc13-1 and PSD-95 nanostructure^78^. Finally, the presynaptic adhesion protein PTP0 also promotes NMDAR responses independent of its trans-synaptic adhesion domain, suggesting it could have a role in regulating release specifically near NMDARs^79^. Future work will need to take advantage of multiplexed imaging and extend acute manipulation techniques^14,80^ to tease apart the complex functional roles of these diverse molecules in regulating NMDAR responses.

We observed significant, though partial, co-enrichment of GluN2A and GluN2B subunit nanodomains, and their selective enrichment with a subset of release sites in the nanocolumn. Previous work has observed minimal overlap between subunit nanodomains^22,23^. This may arise from differences in culture age, expression levels, surface vs total staining, or the imaging modality, though we note that we accomplished our mapping without subunit overexpression.

Nevertheless, the key insight we add is that we have mapped the overlapping GluN2 nanodomains simultaneously and in the molecular context of two key synaptic proteins, which revealed their enrichment to the nanocolumn and suggests their relative importance in the synapse, regardless of whether they are the majority receptor population. This subunit co-enrichment specifically across from nanocolumnar release sites suggests particular synaptic subregions (nanodomains) may facilitate activation of specific NMDAR subtypes. This in turn may facilitate transduction of sparse signals. Ca^2+^ is quickly buffered by calmodulin after entry into the dendritic spine^81,82^, and as only a few of the estimated 10-20 NMDARs per synapse are activated per stimulus^67^, there may also be postsynaptic hotspots of Ca^2+^ influx. Beyond Ca^2+^-dependent signaling, NMDARs also pass non-ionotropic signals via effector proteins such as PP1 that interact directly with the receptor C-tail^18^. Therefore, positioning NMDAR signaling partners within the proposed NMDAR functional nanodomains could facilitate their downstream activation.

Co-enriched GluN2 subunit nanodomains could be constructed from either mixed populations of diheteromers or of triheteromers, which could confer unique signaling properties to these nanodomains. Triheteromers make up ∼50% of synaptic NMDARs in mature synapses^45,46^ and have unique kinetic properties compared to diheteromers^83,84^. Although delineating the trafficking of triheteromers has not yet been feasible, it is notable that a triheteromer carrying dual GluN2 C-tails likely engages in a broader range of interactions than either diheteromer carrying only one type. We observed a slightly higher enrichment of nanocolumnar release sites with GluN2A vs GluN2B, which could come about due to preferential binding of GluN2A over GluN2B to PSD-95^19,85^. However, there is still a significant portion of GluN2A outside the nanocolumn, suggesting other mechanisms are at play. For example, a receptor carrying GluN2A and GluN2B C-tails together could create an avidity effect that increases the range of conformational possibilities with PSD-95, or perhaps the combination of GluN2A and GluN2B binding with other MAGUKs, such as PSD-93 or SAP102, could create a binding environment that gathers both subunits. This could have implications for LTP, as triheteromers have fast, GluN2A-like kinetics and could bring the receptor to PSD-95 nanodomains, while the GluN2B C-tail recruits interactors required for LTP like CaMKII^20^. In fact, when GluN2B diheteromers, but not triheteromers, are blocked, LTP remains intact, but a complete GluN2B subunit deletion ablates LTP^86,87^, suggesting a specific role of triheteromers that might be facilitated by their position relative to release sites. This suggests a nanoscale signaling complex where the very precise spatial combination of the NMDAR coincidence detection mechanism that gates Ca^2+^ influx is combined with the ability to interact with downstream LTP effectors and concentrated near high P_r_ release sites. In the future, direct visualization of NMDARs of specific molecular compositions in the context of release sites and other proteins will help clarify the specific roles each receptor subtype plays at the synapse in neurotransmission and plasticity.

Our observations of NMDAR molecular context we believe help illuminate general rules by which synapses are assembled. We suggest that a critical level at which synaptic function is established is through assembly of specific nanodomain configurations from available cell type-specific components, rather than from following an overall synapse-wide scheme such as a center-surround architecture. Several observations support this idea. Here, we document variability in molecular characteristics of presynaptic release sites within a single active zone and show that NMDAR subsynaptic distribution is dependent on highly local transcellular context. Other recent work has shown that postsynapses contain diverse scaffold molecules beyond PSD-95 that are organized in unique and developmentally regulated NCs^10,88,89^.

Synaptic nanoclustering and trans-synaptic alignment are conserved in evolution and observed across several synapse types^48,90,91^, the detailed characteristics of which depend on cleft-resident synaptic organizing complexes^1,9,11,14^. Further, the specific complement of these proteins differs across cell types and may individualize the nano-organization even of the same proteins at different excitatory synapse types^48^, or related ones at inhibitory synapses^90^. The ability to assemble a range of nanoscale protein relationships substantially broadens the functional range of a synapse. Given that assembly and regulation of multi-protein ensembles at the nanoscale level is a ubiquitous requirement for diverse cell functions, the power of DNA-PAINT super-resolution microscopy to provide high-resolution multiplexed protein localization, along with realistic functional modeling based on super-resolution maps, will be critical for analysis of how conditional distribution features sculpt these complex relationships.

## METHODS

### DNA constructs

pORANGE GFP-Grin2b KI (Addgene plasmid #131487), pFUGW spCas9 (Addgene plasmid #131506) and pFUGW mCherry-KASH (Addgene plasmid #131505) were gifts from Harold MacGillavry. psPAX2 (Addgene plasmid #12260) and pMD2.G (Addgene plasmid #12259) were gifts from Didier Trono. LentiCRISPRv2GFP (LCV2) was a gift from David Feldser (Addgene plasmid #82416). GFP-LRRTM2 knockdown/rescue used for determining anti-EGFP antibody dilution was previously described^14^. SEP-GluN2A and SEP-GluN2B were gifts of Andres Barria. Rat GluN1-1a pcDNA3.1+ was a gift of Gabriela Popescu. pFUGW ORANGE GFP-Grin2b KI was made by subcloning the U6 promoter-sgRNA-GFP-Grin2b donor cassette from pORANGE into the PacI restriction site of pFUGW mCherry-KASH with NEBuilder HiFi assembly, then subsequently removing mCherry-KASH with NEB Q5 site-directed mutagenesis. pFSW myr(Fyn)-EGFP-LDLRct^92^ was made by subcloning a synthetic double-stranded DNA fragment of the promoter and ORF (Twist Bioscience) into the PacI and XbaI sites of pFW (pFUGW with the ubiquitin promoter-EGFP removed by NEB Q5 mutagenesis) with restriction/ligation. LCV2 Grin1 KO EGFP, expressing a validated gRNA targeting GRIN1^93^, was previously described^78^. pEGFP-N1 is from Clontech. DNA constructs are detailed Supplementary Table 1.

### Lentivirus

HEK293T cells (ATCC CRL-3216) were maintained in DMEM + 10% FBS and penicillin/streptomycin at 37°C and 5% CO_2_. For lentiviral production, cells were plated at 5×10^6^ cells/10 cm plate and transfected 12-24h later with 6 µg of either pFUGW ORANGE GFP- Grin2B KI, pFUGW spCas9 or LCV2 Grin1 KO EGFP + 4 µg psPAX2 + 2 µg pMD2.G using PEI for 4-6 hours. After 48h, the media was harvested, debris removed by centrifugation at 1000 RPM for 5 min and 0.45 µm PES filtering, and single use aliquots were frozen at-80°C for long term storage without further concentration. Titers were ∼10^5^ IFU/mL and routinely infected 90% or more of the cells on the coverslip at the volumes used.

### Neuron and HEK culture

All animal procedures were approved by the University of Maryland Animal Use and Care committee. Dissociated hippocampal cultures were prepared from E18 Sprague-Dawley rats of both sexes as described previously^48^ and plated on poly-L-lysine-coated coverslips (#1.5, 18 mm, Warner) at a density of 30,000 cells/coverslip. For most experiments, neurons were infected with 100-150 µl each of pFUGW ORANGE GFP-Grin2b KI and pFUGW spCas9 lentivirus at DIV4-6 and fixed at DIV20-21. For DNA-PAINT of dendritic spines, neurons were transfected with 1 µg of pFSW myr(Fyn)-EGFP-LDLRct at DIV14-16 with Lipofectamine 2000 per manufacturer’s instructions, and fixed at DIV20-21. Neurons were similarly transfected with GFP-LRRTM2 for testing anti-EGFP dilution series or with cell-fill EGFP for testing NMDA- induced permeabilization. For testing anti-GluN2A specificity, neurons were infected with 100 µl LCV2 Grin1 KO EGFP lentivirus at DIV5 and fixed at DIV21. For SEP-GluN2 overexpression tests, HEK cells were plated on poly-L-lysine and fibronectin-coated 18 mm coverslips at a density of 100,000 cells/coverslips, transfected 24h later with 250 ng SEP-GluN2A or GluN2B + 125 ng GluN1-1a + 125 ng mCherry-C1 with Lipofectamine 2000, then maintained for 24h in fresh media with 150 µM APV + 11.25 µM MK-801 before fixation.

### Antibody conjugation and preincubation

Primary antibodies are detailed in Supplementary Table 2, and secondary reagents in Supplementary Table 3. Donkey anti-rabbit IgG was conjugated with Cy3B as previously described^48^. Secondary single domain antibodies (sdAbs) for DNA-PAINT were custom-made by Massive Photonics and each carried one of four oligonucleotide docking strands optimized for DNA-PAINT^37^. To stain multiple targets in the same sample with antibodies from the same species, we preincubated^31,32,48,63,88^ primary antibodies with 2.5-fold molar excess of the appropriate species secondary sdAb labeled with DNA-PAINT docking strands for 20 minutes at room temperature (RT) in either PBS + 100 mM glycine (PBS/Gly) for fixed staining or ACSF (10 mM HEPES pH7.4, 139 mM NaCl, 2.5 mM KCl, 10 mM glucose, 2 mM MgCl_2_, 2 mM CaCl_2_) for live staining, then bound excess sdAb by adding 2-fold molar excess of Fc fragment of the species targeted by the secondary sdAb for a further 20 minutes at RT. Preincubated antibodies for a given incubation were then pooled together and diluted to their final concentrations in PBS/Gly or ACSF for use in immunostaining.

### Immunostaining

For DNA-PAINT of EGFP-Grin2b KIs, KI lentivirus-infected DIV20-21 neurons were removed from culture media and placed in ACSF containing primary antibodies preincubated with sdAbs conjugated to DNA-PAINT docking strands. Neurons were incubated in the antibody mixture at 16°C for 60 min and then transferred to fixative (2% PFA in 10mM MES (pH6.8), 138mM KCl, 3mM MgCl_2_, 2mM EGTA, 320mM sucrose) for 15 min at room temperature. Following fixation, neurons were washed in PBS/Gly 3 x 5 min at RT, permeabilized with 0.3% Triton X-100 for 20 min at RT, and blocked with 10% donkey serum in PBS/Gly + 0.2% Triton X-100 for 45 min at RT. Neurons were then incubated overnight at 4°C with sdAb-preincubated primary antibodies diluted in 50% blocking buffer. Neurons were washed 3 x 5 min in PBS/Gly, postfixed in PBS containing 4% PFA and 4% sucrose for 15 min at RT, and finally washed 3 x 5 min with PBS/Gly.

Uninfected DIV20-21 cells were used for DNA-PAINT of NMDA-treated cells and processed similar to above, though here neurons were stained for surface receptor after fixation to avoid labeling receptors that could be subsequently internalized by drug treatment. Neither our fixation protocol nor drug treatment permeabilized dendrites or synapses; though some soma and large apical dendrites were permeabilized by the fixation (Supplementary Fig. 5), these regions were easily avoided in imaging. Culture medium was replaced with vehicle (water), NMDA (20 µM final) or NMDA (20 µM) + D,L-APV (100 µM final) diluted in pre-warmed ACSF for 3 minutes, then neurons were immediately fixed and washed as above. Neurons were stained post-fixation for surface epitopes with preincubated antibodies for 1 hour at RT and washed 3 x 5 min with PBS/Gly, after which staining proceeded, starting with permeabilization, as above. After the post-overnight incubation wash, neurons were incubated with Atto488-labeled anti-PSD-95 sdAb to identify synapses for DNA-PAINT, then washed and postfixed as above.

In control confocal and STED experiments, cells were stained similar to above, swapping DNA-bearing reagents for largely non-preincubated fluorescent secondary reagents. For testing antibody dilution and specificity (Supplementary Fig. 1), cells were stained as for EGFP-Grin2b KI DNA-PAINT. For testing permeabilization by fixation or NMDA (Supplementary Fig. 5) and for STED (Supplementary Fig. 3), cells were stained as for NMDA treatment DNA-PAINT. Secondary antibodies or nanobodies were diluted in PBS/Gly and applied for 1h at RT after washing overnight primaries. Cells were washed 3 x 5 min in PBS/Gly before postfixing as above. Coverslips were mounted on slides in Abberior Solid Antifade mountant for STED.

For detailed use of antibodies in each experiment, see Supplementary Table 4.

### Confocal microscopy and analysis

Confocal images were acquired on a Nikon TI2 inverted microscope equipped with an Andor Dragonfly spinning disk confocal, a Plan Apo λD 60x/1.42 NA oil immersion objective, and a Plan Apo 20x/0.75 NA air objective. Excitation light (405/488/561/640) was supplied by an Andor ILE and reflected to the sample through a 405/488/561/638 quadband polychroic (Chroma), and emission light was passed through the confocal unit and appropriate emission filters (ET525/50, ET600/50 (Chroma) or Em01-R442/647 (Semrock)) to a Zyla 4.2+ sCMOS camera (Andor). Neurons were imaged at 50% laser power and 200 ms exposure, and Z-stacks were acquired using a piezo fitted in a Nikon stage. Z-stacks were converted to maximum intensity projections using FIJI^94^. GFP-LRRTM2-transfected cells were used as a convenient model system to test a mouse anti-EGFP dilution series to use on EGFP-Grin2b KIs: GFP-LRRTM2 generates a similar synaptic, surface-expressed EGFP epitope for staining, and there are many more transfected cells than Kis allowing more robust measurements. These images were analyzed with a custom FIJI macro that first background subtracted the 1^st^ percentile pixel intensity from each channel, then thresholded the GFP-LRRTM2 signal to create a mask and measured the mean EGFP and anti-EGFP intensities inside the ROI.

### STED microscopy and analysis

STED images were acquired on an Abberior Facility Line STED microscope equipped with an Olympus IX83 base and UPlanXApo 60x/1.42 NA oil immersion objective, 405, 485, 561, and 640 nm lasers for imaging, a pulsed 775 nm STED laser, and photon-counting APDs for emission detection. We identified EGFP-Grin2b KI neurons with green fluorescence emission then acquired 2-color STED images of GluN2A and EGFP-GluN2B line-by-line using 6 µs pixel dwell time and 15 nm pixel size (pinhole 0.91 AU). Abberior STAR635p dye was excited by the 640 nm laser at 26.4% power and depleted by the STED laser at 35% power, with accumulation of 20 lines. Abberior STAR580 dye was excited by the 561 nm laser at 16.7% power and depleted by the STED laser at 30% power, with accumulation of 20 lines. Corresponding confocal-resolution images were acquired prior to each STED image. Data were analyzed using custom FIJI macros. For each confocal image, a Laplacian of Gaussian filtering with 0.8 smoothing and a Gaussian blurring were applied. Then, a morphological filter was used upon image binarization, and subtraction of the binary masks allowed the identification of ROIs with signal from both GluN2A and EGFP-GluN2B. These ROIs were then used to mask synapses in STED images. Nanoclusters were identified from a Gaussian function fitted around (6 pixels) identified local intensity maxima (visually assessed prominence: 100). The fitting script was adapted from the “FWHM_along_line_v1” FIJI macro created by Dominic Waithe (https://github.com/dwaithe/generalMacros). XY Coordinates of fitted nanoclusters were then used to determine the nearest-neighbor distance from GluN2B to GluN2A for each staining condition. Extreme outliers, likely representing mis-segmented NCs, were removed from both FWHM and nearest-neighbor distributions using GraphPad PRISM’s ROUT method with strictest settings (Q = 0.1%).

### Single-molecule microscopy

DNA-PAINT images of EGFP-Grin2b KIs were acquired on an Olympus IX81 inverted microscope with an Olympus 100x/1.49 NA TIRF oil immersion objective. Excitation light (405/488/561) from an Andor ALC and a Toptica iBeam Smart (640) was reflected to the sample through a 405/488/561/638 quadband polychroic (Chroma) at an incident angle greater than the critical angle to achieve Highly Inclined and Laminated Optical (HILO) illumination. Emission light was passed through an adaptive optics device (MicAO, Imagine Optic), which corrected aberrations present in the point-spread function, followed by a DV2 image splitter (Photometrics) equipped with a T640lpxr dichroic and ET655lp single band (far-red) and 59004m dual band (red and green) emission filters to allow identification of GFP-Grin2b KI cells with the 488 nm laser followed by simultaneous collection of red and far-red emissions during DNA-PAINT imaging. Emission was finally collected on an iXon+ 897 EM-CCD camera (Andor). Z stability was maintained by the Olympus ZDC2 feedback positioning system. The microscope, ALC, and camera were controlled by iQ3 (Andor), the Toptica laser by TOPAS iBeam Smart GUI, and the Micao by separate Imagine Optic software. An additional arc lamp provided epifluorescence illumination for identifying GFP-Grin2b KI cells. The microscope was contained inside an insulated box with temperature control to minimize sample drift.

DNA-PAINT images for the NMDA treatment experiment were acquired on similar microscope we have described recently^63,78^. Briefly, the system was built on a Nikon TI2 base equipped with a 100x/1.49NA Apo TIRF oil immersion objective, with excitation light provided by an Oxxius L6Cc laser combiner with 405, 488, 561, and 640 nm lasers fiber-coupled to a Nikon manual TIRF illuminator; the emission path was the same as the Olympus system. Z-stability was maintained by the Nikon Perfect Focus System, with Nikon Elements software controlling all equipment except the Micao.

90 nm gold nanoparticles (Cytodiagnostics) were added at a 1:3 dilution for 10 minutes before imaging to act as fiducials for drift and chromatic correction. For imaging of EGFP-GluN2B, KI cells were identified based on GFP-Booster AF488 staining and selected to have similar AF488 intensity and cell morphology across experiments, targeting apparently spiny regions with few apparent shaft synapses on spiny, pyramidal-shaped neurons. Four targets were imaged in two exchange rounds. Cy3B and Atto643 DNA-PAINT imager strands (Massive Photonics) (one each) were diluted into imaging buffer (1x PBS pH7.4 + 500 mM NaCl + oxygen scavengers (PCA/PCD/Trolox))^30^ to the indicated concentrations and added to the sample. Drift was allowed to settle for 10 minutes, then 50,000 frames were acquired with 50 ms exposure. Output laser power at the objective on the Olympus setup was ∼27 mW for the 640 nm laser and ∼18 mW for the 561 nm laser, yielding power densities of ∼3.3 and ∼2.2 and kW/cm^2^, respectively. After acquisition, the imagers were removed by gently exchanging the imaging buffer with 20 mL exchange buffer (1x PBS pH7.4), then replacing the exchange buffer with fresh imaging buffer containing the next set of imager strands. After letting drift settle 10 minutes, the second round of imaging was acquired as before. Images were acquired similarly for the NMDA treatment experiment with minor modifications, similar to^63^. Isolated stretches of dendrite with apparent dendritic spines were identified based on PSD-95 nanobody Atto488 staining, and three targets were imaged in two exchange rounds. Cy3B and Atto655 imager strands were diluted into imaging buffer without oxygen scavengers, and 30,000 frames were acquired with 100 ms exposure. Output laser power density at the objective on the Nikon setup was ∼170 W/cm^2^ for the 640 nm laser and ∼87.4 W/cm^2^ for the 561 nm laser. TetraSpeck beads (100 nm; Invitrogen) were immobilized on separate coverslips prepared with poly-L-lysine as for the cultured neurons, and 8-10 fields of beads were imaged for 100 frames at 50 ms exposure each and used to generate transforms to correct chromatic aberrations between the two channels.

### Single molecule processing

Images were processed in batch with a combination of FIJI, Picasso v0.4.11 (https://github.com/jungmannlab/picasso), and custom MATLAB scripts similar to our previous work^48,63,88^. The analysis pipeline is described in detail in Supplementary Note 1 and Supplementary Figure 2. In brief, images were localized and drift corrected in Picasso, and chromatic aberrations between channels corrected with a custom MATLAB script. Localizations were subsequently filtered to remove spurious detections and linked to combine localizations persisting for more than one frame. Clusters of synaptic proteins were identified with DBSCAN and removed if they displayed kinetic properties of non-specific imager binding^95^. Finally, high-confidence synapses were picked by manually inspecting for the presence of the other imaged proteins in sufficient density for analysis and to confirm that the kept cluster was a synapse.

Synapses were kept based on disclike shapes, overlap of pre-and post-synaptic proteins, a size range of ∼100 – 800 nm diameter, and their position near a dendrite, then scored as “en face”, “side view” or “intermediate”, ie, somewhere between en face and side. Synapses were then further filtered for en face by removing those with a long/short axis ratio >2, then validated independently by three expert raters. In some cases, super-resolution images were rendered using the FIJI ThunderSTORM plugin’s^96^ *average shifted histogram* method with 10 nm pixels (magnification 16) for ease of visualization. Otherwise, localizations are plotted as heat maps of local density, calculated for each localization as the number of localizations within 2x epsilon (see below) for that protein and normalizing to the maximum value per synapse.

### Single-molecule analyses

*Synapse analyses:* Protein autocorrelations (AC) and cross-correlations (CC) were determined using custom MATLAB functions as previously described^88,97^ with 5 nm render pixels and a max shift radius of 500 nm for AC and 5 nm render pixels and max shift radius of 250 nm for the CC. NCs were detected using DBSCAN with the following parameters: GluN2B epsilon = 16 nm, minpts = 11; GluN2A epsilon = 16 nm, minpts = 8; Munc13-1 epsilon = 17.6 nm, minpts = 9.

DBSCAN parameters for PSD-95 were set per synapse to normalize for density variations between synapses, where epsilon was 5x standard deviations greater than the mean minimal distance of 50 randomizations of PSD-95 localization positions within the same space, and minpts was 5x standard deviations greater than the mean number of points within that epsilon. NC areas were determined using the MATLAB *alphaShape* function with ‘HoleThreshold’ set to suppress all interior holes and an alpha radius of 150 nm, followed by the *area* function. NCs with fewer than five localizations or with large outlier areas due to erroneous grouping by DBSCAN were removed from analysis (area maximums identified by GraphPad Prism’s ROUT method at Q = 0.1%: GluN2A = 2430.7 µm^2^, GluN2B = 3376.6 µm^2^, PSD-95 = 6202.9 µm^2^, Munc13-1 = 2818.6 µm^2^). To calculate NC position relative to PSD center and edge, PSD-95 localizations at each synapse were fit to an ellipse (modifying an approach by Nima Moshtagh (2007); Minimum Volume Enclosing Ellipsoid v1.2.0.0, https://www.mathworks.com/matlabcentral/fileexchange/9542-minimum-volume-enclosing-ellipsoid, MATLAB Central File Exchange, retrieved May 4, 2023) that was used to derive a concentric ellipse that included the center of a NC of interest on its perimeter. The ratio of the area of these two ellipses yields a two-dimensional measure of how close to the center or to the edge a given NC is located, the square root of which results in a linear representation, where 0 is a NC at the center of the synapse, 1 is a NC on the edge of the synapse, and 0.5 is a NC that is positioned halfway between center and edge. The distance of extrasynaptic NCs to the PSD border was calculated from a vector connecting the extrasynaptic NC and PSD centroids by subtracting the distance between the PSD centroid and border (determined using MATLAB’s *intersect* function) from the distance between the PSD and NC centroids.

*Nanocluster analyses:* NC peak-to-peak distances were determined as the linear distance (MATLAB *pdist2*) from the peak of a NC to the nearest peak of another protein NC. NC pairing was performed by, for each NC, identifying based on peak-to-peak distances whether there existed a NC of a second protein that was mutually nearest to it. NC overlap was determined using the separation index as described previously^63^, which normalizes the distance between NCs to the sum of their radii, resulting in a measure ranging from 0 (perfect overlap) to 1 (perfectly adjacent) and greater (no overlap).

*Cross-enrichments:* Cross-enrichments (CE) were determined as described^63,88,97^. In brief, the peak density of one protein NC was used as the reference point, and the distance of all localizations of a second protein, or of a modified randomized synapse of that protein with 300x more localizations to avoid bins with zero localizations^48,63^, were determined to this point. CE is calculated as the number of localizations with distances to the reference position in 10 nm bins normalized to the same for the randomized synapse, with the 300x density factor divided out.

CEs can be noisy due to the randomization resulting in a value that disproportionately represents the true density within a distance bin. To retain the information in that bin but minimize this noisiness, CEs were smoothed by using the MATLAB *isoutlier* function and its ‘quartiles’ method to detect outliers per distance bin and replacing the outlier values with the largest, non-outlier value in that bin. Auto-enrichments (AE) were calculated the same way except the distance of localizations of one protein was measured in reference to the peak density of its own NC. NC density was calculated as the mean of the first 60 nm of the auto-enrichment.

*Conditional comparisons:* Nanoclusters were determined to be “enriched” or “not enriched” if the enrichment index (average of the first 60 nm of the CE) to the real data was greater than or less than 1.96x standard deviations from the mean of the enrichment index to a mean randomized synapse, respectively. Nanoclusters were then subset into these groups and their CE with another protein plotted. For example, a Munc13-1 NC can be conditioned on its nanocolumn status, ie, within the nanocolumn (Munc13-1 NC enriched with PSD-95) or outside the nanocolumn (Munc13-1 NC de-enriched with PSD-95), then the CE of those Munc13-1 NC groups with GluN2 subunits compared. To calculate the AE and CE of GluN2 subunits subset by nanocolumn status, we first identified the nearest GluN2 NC to each Munc13-1 NC by mutually closest pairing as described above (removing any mutual pair greater than 60 nm apart), then subset the GluN2 NCs by their paired Munc13-1 nanocolumn status.

### MCell modeling and analysis

We modeled the magnitude and time course of NMDAR activation in response to glutamate release using MCell^49,50^. MCell allows realistic diffusion-reaction simulation using complex spatial models; each receptor type, release position, and glutamate molecule, as well as their interactions, can thus be tracked individually, allowing determination of the functional impact of release site position on receptor activation. We used experimental results from a subset of super-resolved synapses to constrain the placement of biomolecules and synapse structures for MCell models, and constructed 3D geometries of synapses and their borders using Blender (https://www.blender.org/) and CellBlender (https://mcell.org/). The synapses selected for analysis were required to have both nanocolumnar and non-nanocolumnar release sites for both GluN2A and GluN2B NCs to permit within-synapse comparisons. We defined release site positions using Munc13-1 NC peaks, GluN2 subunit positions using their localizations, and the bounds of the synapse using a minimal ellipse containing the combined PSD-95, GluN2A, GluN2B, and Munc13-1 localizations. For each of the modeled synapses, we ran three types of simulations to compare the receptor activation probability for the set of experimentally obtained release sites and receptor positions versus uniformly distributed release sites or receptor positions: 1) real positions of release sites and receptors, 2) real positions of receptors and uniform positions of release sites, 3) real positions of release sites and uniform positions of receptors. Uniformly distributed positions were created as a grid of points with 50 nm spacing in x and y and located inside the synapse border (ellipse).

We modeled the NMDAR kinetic schemes to predict the opening rates of each receptor in response to glutamate release based on the kinetic schemes derived from Santucci et al.^47^, and used an analytical model^98^ for release of glutamate in the synaptic cleft. Glutamate molecules that reached the edge of the cleft volume disappeared from the simulation run. For computational efficiency we approximated glutamate release through vesicle fusion by emitting glutamate from the vesicle at the correct rate with Poisson probabilities computed from a time-dependent rate constant k(t)^99,100^. We converted the number of glutamate molecules released to the time dependent rate of release, and the time-dependent rate constant k was precomputed and stored in separate text files that were then read by MCell at the beginning of each simulation. We ran 50 independent simulations using different random numbers (seeds) and tracked all the reactions in the system and measured the probability of receptor opening. The simulation results were then analyzed using in-house scripts on Bridges-2 machines at the Pittsburgh Supercomputing Center using regular memory nodes (256 gigabytes per node). We used MDL user interface to design and control MCell version 3.4; key simulation parameters are included in Supplementary Table 6. In total about 1.6 million simulations were run and analyzed.

The open fraction of NMDARs (P_o_) for each subunit localization was calculated by dividing the number of trials where the receptor opened by the total number of trials at each time step; for total NMDAR P_o_, the number of open trials for both subunits was summed at each timepoint then divided by the summed number of trials. To allow determination of peak P_o_ even from noisy traces or those without a clear peak, the P_o_ time course of all localizations that showed receptor opening was averaged and a small time window identified around the peak of this trace (peak P_o_ window: 2.5 +/-1 ms for GluN2A, 5.5 +/- 2 ms for GluN2B). The peak P_o_ for each subunit in each simulation run was then calculated as the average P_o_ within this time window. Synaptic P_o_ was calculated as the average peak P_o_ of all subunit localizations for each synapse. Data were grouped by release site, nanocluster, or nanocolumn status as indicated.

### Statistics

For experiments imaging EGFP-Grin2b KIs, 74 high-confidence, en face synapses from 8 neurons over 6 independent cultures were measured. Glutamate release modeling was performed using super-resolution maps from 8 representative synapses in this dataset. For the NMDA treatment experiment, 76 vehicle, 80 NMDA, and 82 NMDA+APV-treated synapses from 8 neurons each over 4 independent cultures were measured, and the experimenters were blinded to condition throughout imaging and analysis. For STED imaging, 1504 GluN2A and 1403 EGFP-GluN2B NCs (1°Nb staining group), and 2043 GluN2A and 1775 EGFP-GluN2B NCs (1°Ab + 2°Nb staining group), from 7 and 6 neurons, respectively, over 2 independent cultures were measured, and the experimenters were blinded to condition throughout imaging and analysis. All statistical comparisons were performed in GraphPad Prism 10. Data were tested for normality using a Kolmogorov-Smirnov test. All normally distributed datasets in this work met the assumptions of homoscedasticity (F test), and differences between groups were tested using two-tailed unpaired t-tests. For non-parametric data, differences between groups were tested using two-tailed Mann-Whitney tests for two groups or a Kruskal-Wallis test, followed by post-hoc Dunn’s multiple comparisons tests, for greater than two groups.

Differences between groups in modeling data were tested using two-tailed paired t-test.

## Data availability

All analyzed data are included in the figures and will be available at publication on the Open Science Framework at https://osf.io/5wvhk/. Original data generated in the current study are available from the corresponding authors on reasonable request. Source code for analyses will be made available at publication at https://github.com/blanpiedlab.

## Author contributions

The project was conceptualized by MCA, PAD, SRM, TAB, and ADL. Experimental conditions and reagents were optimized by MCA, MD, and ADL. Confocal data were collected by MCA and ADL. STED data were collected and analyzed by MCA and MD. DNA-PAINT data were collected by ADL and analysis was performed by MCA, ADL, PAD, and SRM. Modeling data were generated by RL and analyzed by RL and ADL. TAB and ADL supervised the study. Figures were prepared by MCA and ADL, and the manuscript was written by ADL with constructive review and editing by all authors.

## Acknowledgements

This work was supported by funds from the National Institutes of Health (F31MH124283 to MCA, F31MH117920 to PAD, R37MH080046 and R01MH11982 to TAB, and F32MH119687 to ADL). STED images were acquired in the University of Maryland School of Medicine Center for Innovative Biomedical Resources Core Confocal Facility, with support from the University of Maryland Medicine Institute for Neuroscience Discovery (UM-MIND). This work used the Extreme Science and Engineering Discovery Environment (XSEDE), which is supported by the National Science Foundation (ACI-1548562). Specifically, it used the Bridges-2 system, which is supported by the National Science Foundation (ACI-1928147), at the Pittsburgh Supercomputing Center (PSC). We thank Minerva Contreras for excellent technical support, Dr. Shilpa Dilip Kumar for assistance on STED imaging, and the Blanpied lab for helpful discussions.

## REFERENCES

1. Fukata, Y. et al. LGI1-ADAM22-MAGUK configures transsynaptic nanoalignment for synaptic transmission and epilepsy prevention. Proc. Natl. Acad. Sci. U. S. A. 118, e2022580118 (2021).

2. Zhu, W.-H. et al. Nanoscale reorganisation of synaptic proteins in Alzheimer’s disease. Neuropathol. Appl. Neurobiol. 49, e12924 (2023).

3. Zieger, H. L. & Choquet, D. Nanoscale synapse organization and dysfunction in neurodevelopmental disorders. Neurobiol. Dis. 158, 105453 (2021).

4. Choquet, D. & Opazo, P. The role of AMPAR lateral diffusion in memory. Semin. Cell Dev. Biol. 125, 76–83 (2022).

5. Eggermann, E., Bucurenciu, I., Goswami, S. P. & Jonas, P. Nanodomain coupling between Ca^2+^ channels and sensors of exocytosis at fast mammalian synapses. Nat. Rev. Neurosci. 13, 7–21 (2011).

6. Gou, X.-Z., Ramsey, A. M. & Tang, A.-H. Re-examination of the determinants of synaptic strength from the perspective of superresolution imaging. Curr. Opin. Neurobiol. 74, 102540 (2022).

7. Guzikowski, N. J. & Kavalali, E. T. Nano-Organization at the Synapse: Segregation of Distinct Forms of Neurotransmission. Front. Synaptic Neurosci. 13, 796498 (2021).

8. Biederer, T., Kaeser, P. S. & Blanpied, T. A. Transcellular Nanoalignment of Synaptic Function. Neuron 96, 680–696 (2017).

9. Lloyd, B. A., Han, Y., Roth, R., Zhang, B. & Aoto, J. Neurexin-3 subsynaptic densities are spatially distinct from Neurexin-1 and essential for excitatory synapse nanoscale organization in the hippocampus. Nat. Commun. 14, 4706 (2023).

10. Tang, A.-H. et al. A trans-synaptic nanocolumn aligns neurotransmitter release to receptors. Nature 536, 210 (2016).

11. Haas, K. T. et al. Pre-post synaptic alignment through neuroligin-1 tunes synaptic transmission efficiency. eLife 7, e31755 (2018).

12. Han, Y. et al. Neuroligin-3 confines AMPA receptors into nanoclusters, thereby controlling synaptic strength at the calyx of Held synapses. Sci. Adv. 8, eabo4173 (2022).

13. Sinnen, B. L. et al. Optogenetic Control of Synaptic Composition and Function. Neuron 93, 646–660.e5 (2017).

14. Ramsey, A. M. et al. Subsynaptic positioning of AMPARs by LRRTM2 controls synaptic strength. Sci. Adv. 7, eabf3126 (2021).

15. Paoletti, P., Bellone, C. & Zhou, Q. NMDA receptor subunit diversity: impact on receptor properties, synaptic plasticity and disease. Nat. Rev. Neurosci. 14, 383–400 (2013).

16. Hansen, K. B., Yi, F., Perszyk, R. E., Menniti, F. S. & Traynelis, S. F. NMDA Receptors in the Central Nervous System. in NMDA Receptors (eds. Burnashev, N. & Szepetowski, P.) vol. 1677 1–80 (Springer New York, New York, NY, 2017).

17. Hosokawa, T. & Liu, P.-W. Regulation of the Stability and Localization of Post-synaptic Membrane Proteins by Liquid-Liquid Phase Separation. Front. Physiol. 12, 795757 (2021).

18. Dore, K. et al. Unconventional NMDA Receptor Signaling. J. Neurosci. Off. J. Soc. Neurosci. 37, 10800–10807 (2017).

19. Gardoni, F. & Di Luca, M. Protein-protein interactions at the NMDA receptor complex: From synaptic retention to synaptonuclear protein messengers. Neuropharmacology 190, 108551 (2021).

20. Hardingham, G. NMDA receptor C-terminal signaling in development, plasticity, and disease. F1000Research 8, F1000 Faculty Rev-1547 (2019).

21. Petit-Pedrol, M. & Groc, L. Regulation of membrane NMDA receptors by dynamics and protein interactions. J. Cell Biol. 220, e202006101 (2021).

22. Hruska, M., Cain, R. E. & Dalva, M. B. Nanoscale rules governing the organization of glutamate receptors in spine synapses are subunit specific. Nat. Commun. 13, 920 (2022).

23. Kellermayer, B. et al. Differential Nanoscale Topography and Functional Role of GluN2- NMDA Receptor Subtypes at Glutamatergic Synapses. Neuron 100, 106–119.e7 (2018).

24. Sun, Y. et al. The differences between GluN2A and GluN2B signaling in the brain. J. Neurosci. Res. 96, 1430–1443 (2018).

25. Niethammer, M., Kim, E. & Sheng, M. Interaction between the C terminus of NMDA receptor subunits and multiple members of the PSD-95 family of membrane-associated guanylate kinases. J. Neurosci. Off. J. Soc. Neurosci. 16, 2157–2163 (1996).

26. Böhme, M. A. et al. Active zone scaffolds differentially accumulate Unc13 isoforms to tune Ca(2+) channel-vesicle coupling. Nat. Neurosci. 19, 1311–1320 (2016).

27. Midorikawa, M., Sakamoto, H., Nakamura, Y., Hirose, K. & Miyata, M. Developmental refinement of the active zone nanotopography and axon wiring at the somatosensory thalamus. Cell Rep. 43, 114770 (2024).

28. Reddy-Alla, S. et al. Stable Positioning of Unc13 Restricts Synaptic Vesicle Fusion to Defined Release Sites to Promote Synchronous Neurotransmission. Neuron 95, 1350–1364.e12 (2017).

29. Sakamoto, H. et al. Synaptic weight set by Munc13-1 supramolecular assemblies. Nat. Neurosci. 21, 41–49 (2018).

30. Schnitzbauer, J., Strauss, M. T., Schlichthaerle, T., Schueder, F. & Jungmann, R. Super-resolution microscopy with DNA-PAINT. Nat. Protoc. 12, 1198–1228 (2017).

31. Pleiner, T., Bates, M. & Görlich, D. A toolbox of anti-mouse and anti-rabbit IgG secondary nanobodies. J. Cell Biol. 217, 1143–1154 (2018).

32. Sograte-Idrissi, S. et al. Circumvention of common labelling artefacts using secondary nanobodies. Nanoscale 12, 10226–10239 (2020).

33. Willems, J., et al. ORANGE: A CRISPR/Cas9-based genome editing toolbox for epitope tagging of endogenous proteins in neurons. PLoS Biol. 18, e3000665 (2020).

34. Fang, H., Bygrave, A. M., Roth, R. H., Johnson, R. C. & Huganir, R. L. An optimized CRISPR/Cas9 approach for precise genome editing in neurons. eLife 10, e65202 (2021).

35. Luo, J.-H. et al. Functional expression of distinct NMDA channel subunits tagged with green fluorescent protein in hippocampal neurons in culture. Neuropharmacology 42, 306– 318 (2002).

36. Friedl, K. et al. Assessing crosstalk in simultaneous multicolor single-molecule localization microscopy. *Cell Rep*. Methods 3, 100571 (2023).

37. Strauss, S. & Jungmann, R. Up to 100-fold speed-up and multiplexing in optimized DNA-PAINT. Nat. Methods 17, 789–791 (2020).

38. Dupuis, J. P. et al. Surface dynamics of GluN2B-NMDA receptors controls plasticity of maturing glutamate synapses. EMBO J. 33, 842–861 (2014).

39. Broadhead, M. J. et al. PSD95 nanoclusters are postsynaptic building blocks in hippocampus circuits. Sci. Rep. 6, 24626 (2016).

40. Gürth, C.-M. et al. Neuronal activity modulates the incorporation of newly translated PSD- 95 into a robust structure as revealed by STED and MINFLUX. BioRxiv Prepr. Serv. Biol. (2023) doi:10.1101/2023.10.18.562700.

41. MacGillavry, H. D., Song, Y., Raghavachari, S. & Blanpied, T. A. Nanoscale scaffolding domains within the postsynaptic density concentrate synaptic AMPA receptors. Neuron 78, 615–622 (2013).

42. Khater, I. M., Nabi, I. R. & Hamarneh, G. A Review of Super-Resolution Single-Molecule Localization Microscopy Cluster Analysis and Quantification Methods. Patterns N. Y. N 1, 100038 (2020).

43. Dani, A., Huang, B., Bergan, J., Dulac, C. & Zhuang, X. Superresolution imaging of chemical synapses in the brain. Neuron 68, 843–856 (2010).

44. Kortus, S. et al. Subunit-Dependent Surface Mobility and Localization of NMDA Receptors in Hippocampal Neurons Measured Using Nanobody Probes. J. Neurosci. Off. J. Soc. Neurosci. 43, 4755–4774 (2023).

45. Al-Hallaq, R. A., Conrads, T. P., Veenstra, T. D. & Wenthold, R. J. NMDA di-heteromeric receptor populations and associated proteins in rat hippocampus. J. Neurosci. Off. J. Soc. Neurosci. 27, 8334–8343 (2007).

46. Rauner, C. & Köhr, G. Triheteromeric NR1/NR2A/NR2B receptors constitute the major N-methyl-D-aspartate receptor population in adult hippocampal synapses. J. Biol. Chem. 286, 7558–7566 (2011).

47. Santucci, D. M. & Raghavachari, S. The effects of NR2 subunit-dependent NMDA receptor kinetics on synaptic transmission and CaMKII activation. PLoS Comput. Biol. 4, e1000208 (2008).

48. Dharmasri, P. A., Levy, A. D. & Blanpied, T. A. Differential nanoscale organization of excitatory synapses onto excitatory vs. inhibitory neurons. Proc. Natl. Acad. Sci. U. S. A. 121, e2315379121 (2024).

49. Kerr, R. A. et al. FAST MONTE CARLO SIMULATION METHODS FOR BIOLOGICAL REACTION-DIFFUSION SYSTEMS IN SOLUTION AND ON SURFACES. SIAM J. Sci. Comput. Publ. Soc. Ind. Appl. Math. 30, 3126 (2008).

50. Stiles, J. R., Van Helden, D., Bartol, T. M., Salpeter, E. E. & Salpeter, M. M. Miniature endplate current rise times less than 100 microseconds from improved dual recordings can be modeled with passive acetylcholine diffusion from a synaptic vesicle. Proc. Natl. Acad. Sci. U. S. A. 93, 5747–5752 (1996).

51. Lee, H. K., Kameyama, K., Huganir, R. L. & Bear, M. F. NMDA induces long-term synaptic depression and dephosphorylation of the GluR1 subunit of AMPA receptors in hippocampus. Neuron 21, 1151–1162 (1998).

52. Kaeser, P. S. & Regehr, W. G. Molecular mechanisms for synchronous, asynchronous, and spontaneous neurotransmitter release. Annu. Rev. Physiol. 76, 333–363 (2014).

53. Kusick, G. F. et al. Synaptic vesicles transiently dock to refill release sites. Nat. Neurosci. 23, 1329–1338 (2020).

54. Melom, J. E., Akbergenova, Y., Gavornik, J. P. & Littleton, J. T. Spontaneous and evoked release are independently regulated at individual active zones. J. Neurosci. Off. J. Soc. Neurosci. 33, 17253–17263 (2013).

55. Mendonça, P. R. F. et al. Asynchronous glutamate release is enhanced in low release efficacy synapses and dispersed across the active zone. Nat. Commun. 13, 3497 (2022).

56. Wang, C. S., Chanaday, N. L., Monteggia, L. M. & Kavalali, E. T. Probing the segregation of evoked and spontaneous neurotransmission via photobleaching and recovery of a fluorescent glutamate sensor. eLife 11, e76008 (2022).

57. Atasoy, D. et al. Spontaneous and evoked glutamate release activates two populations of NMDA receptors with limited overlap. J. Neurosci. Off. J. Soc. Neurosci. 28, 10151–10166 (2008).

58. Reese, A. L. & Kavalali, E. T. Single synapse evaluation of the postsynaptic NMDA receptors targeted by evoked and spontaneous neurotransmission. eLife 5, e21170 (2016).

59. Sutton, M. A., Wall, N. R., Aakalu, G. N. & Schuman, E. M. Regulation of dendritic protein synthesis by miniature synaptic events. Science 304, 1979–1983 (2004).

60. Sutton, M. A., Taylor, A. M., Ito, H. T., Pham, A. & Schuman, E. M. Postsynaptic decoding of neural activity: eEF2 as a biochemical sensor coupling miniature synaptic transmission to local protein synthesis. Neuron 55, 648–661 (2007).

61. Li, S. et al. Asynchronous release sites align with NMDA receptors in mouse hippocampal synapses. Nat. Commun. 12, 677 (2021).

62. Karlocai, M. R. et al. Variability in the Munc13-1 content of excitatory release sites. eLife 10, e67468 (2021).

63. Emperador-Melero, J. et al. Distinct active zone protein machineries mediate Ca2+ channel clustering and vesicle priming at hippocampal synapses. Nat. Neurosci. 1–15 (2024) doi:10.1038/s41593-024-01720-5.

64. Maschi, D. & Klyachko, V. A. Spatiotemporal dynamics of multi-vesicular release is determined by heterogeneity of release sites within central synapses. eLife 9, e55210 (2020).

65. Aldahabi, M. et al. Different priming states of synaptic vesicles underlie distinct release probabilities at hippocampal excitatory synapses. Neuron 110, 4144–4161.e7 (2022).

66. Goncalves, J. et al. Nanoscale co-organization and coactivation of AMPAR, NMDAR, and mGluR at excitatory synapses. Proc. Natl. Acad. Sci. U. S. A. 117, 14503–14511 (2020).

67. Nimchinsky, E. A., Yasuda, R., Oertner, T. G. & Svoboda, K. The number of glutamate receptors opened by synaptic stimulation in single hippocampal spines. J. Neurosci. Off. J. Soc. Neurosci. 24, 2054–2064 (2004).

68. Fan, X., Jin, W. Y. & Wang, Y. T. The NMDA receptor complex: a multifunctional machine at the glutamatergic synapse. Front. Cell. Neurosci. 8, 160 (2014).

69. Myers, J. B. et al. The CaMKII holoenzyme structure in activation-competent conformations. Nat. Commun. 8, 15742 (2017).

70. Raghavachari, S. & Lisman, J. E. Properties of quantal transmission at CA1 synapses. J. Neurophysiol. 92, 2456–2467 (2004).

71. Lisman, J., Yasuda, R. & Raghavachari, S. Mechanisms of CaMKII action in long-term potentiation. Nat. Rev. Neurosci. 13, 169–182 (2012).

72. Hosokawa, T. et al. CaMKII activation persistently segregates postsynaptic proteins via liquid phase separation. Nat. Neurosci. 24, 777–785 (2021).

73. Warnet, X. L., Bakke Krog, H., Sevillano-Quispe, O. G., Poulsen, H. & Kjaergaard, M. The C-terminal domains of the NMDA receptor: How intrinsically disordered tails affect signalling, plasticity and disease. Eur. J. Neurosci. 54, 6713–6739 (2021).

74. Jang, S., Lee, H. & Kim, E. Synaptic adhesion molecules and excitatory synaptic transmission. Curr. Opin. Neurobiol. 45, 45–50 (2017).

75. Dai, J., Aoto, J. & Südhof, T. C. Alternative Splicing of Presynaptic Neurexins Differentially Controls Postsynaptic NMDA and AMPA Receptor Responses. Neuron 102, 993–1008.e5 (2019).

76. Budreck, E. C. et al. Neuroligin-1 controls synaptic abundance of NMDA-type glutamate receptors through extracellular coupling. Proc. Natl. Acad. Sci. U. S. A. 110, 725–730 (2013).

77. Nolt, M. J. et al. EphB controls NMDA receptor function and synaptic targeting in a subunit-specific manner. J. Neurosci. Off. J. Soc. Neurosci. 31, 5353–5364 (2011).

78. Dharmasri, P. A., DeMarco, E. M., Anderson, M. C., Levy, A. D. & Blanpied, T. A. Loss of postsynaptic NMDARs drives nanoscale reorganization of Munc13-1 and PSD-95. BioRxiv Prepr. Serv. Biol. 2024.01.12.574705 (2024) doi:10.1101/2024.01.12.574705.

79. Kim, K. et al. Presynaptic PTPσ regulates postsynaptic NMDA receptor function through direct adhesion-independent mechanisms. eLife 9, e54224 (2020).

80. Xu, N. et al. Structural and functional reorganization of inhibitory synapses by activity-dependent cleavage of neuroligin-2. Proc. Natl. Acad. Sci. U. S. A. 121, e2314541121 (2024).

81. Faas, G. C., Raghavachari, S., Lisman, J. E. & Mody, I. Calmodulin as a direct detector of Ca2+ signals. Nat. Neurosci. 14, 301–304 (2011).

82. Sabatini, B. L., Oertner, T. G. & Svoboda, K. The life cycle of Ca(2+) ions in dendritic spines. Neuron 33, 439–452 (2002).

83. Hansen, K. B., Ogden, K. K., Yuan, H. & Traynelis, S. F. Distinct functional and pharmacological properties of Triheteromeric GluN1/GluN2A/GluN2B NMDA receptors. Neuron 81, 1084–1096 (2014).

84. Stroebel, D., Carvalho, S., Grand, T., Zhu, S. & Paoletti, P. Controlling NMDA receptor subunit composition using ectopic retention signals. J. Neurosci. Off. J. Soc. Neurosci. 34, 16630–16636 (2014).

85. Sans, N. et al. A developmental change in NMDA receptor-associated proteins at hippocampal synapses. J. Neurosci. Off. J. Soc. Neurosci. 20, 1260–1271 (2000).

86. Delaney, A. J., Sedlak, P. L., Autuori, E., Power, J. M. & Sah, P. Synaptic NMDA receptors in basolateral amygdala principal neurons are triheteromeric proteins: physiological role of GluN2B subunits. J. Neurophysiol. 109, 1391–1402 (2013).

87. Foster, K. A. et al. Distinct roles of NR2A and NR2B cytoplasmic tails in long-term potentiation. J. Neurosci. Off. J. Soc. Neurosci. 30, 2676–2685 (2010).

88. Metzbower, S. R., Levy, A. D., Dharmasri, P. A., Anderson, M. C. & Blanpied, T. A. Distinct SAP102 and PSD-95 nano-organization defines multiple types of synaptic scaffold protein domains at single synapses. J. Neurosci. Off. J. Soc. Neurosci. e1715232024 (2024) doi:10.1523/JNEUROSCI.1715-23.2024.

89. Sun, S.-Y. et al. Correlative Assembly of Subsynaptic Nanoscale Organizations During Development. Front. Synaptic Neurosci. 14, 748184 (2022).

90. Crosby, K. C. et al. Nanoscale Subsynaptic Domains Underlie the Organization of the Inhibitory Synapse. Cell Rep. 26, 3284–3297.e3 (2019).

91. Muttathukunnel, P., Frei, P., Perry, S., Dickman, D. & Müller, M. Rapid homeostatic modulation of transsynaptic nanocolumn rings. Proc. Natl. Acad. Sci. U. S. A. 119, e2119044119 (2022).

92. Kameda, H. et al. Targeting green fluorescent protein to dendritic membrane in central neurons. Neurosci. Res. 61, 79–91 (2008).

93. Incontro, S., Asensio, C. S., Edwards, R. H. & Nicoll, R. A. Efficient, complete deletion of synaptic proteins using CRISPR. Neuron 83, 1051–1057 (2014).

94. Schindelin, J., et al. Fiji: an open-source platform for biological-image analysis. Nat. Methods 9, 676–682 (2012).

95. Sun, C. et al. The prevalence and specificity of local protein synthesis during neuronal synaptic plasticity. Sci. Adv. 7, eabj0790 (2021).

96. Ovesný, M., Křížek, P., Borkovec, J., Svindrych, Z. & Hagen, G. M. ThunderSTORM: a comprehensive ImageJ plug-in for PALM and STORM data analysis and super-resolution imaging. Bioinforma. Oxf. Engl. 30, 2389–2390 (2014).

97. Chen, J.-H., Blanpied, T. A. & Tang, A.-H. Quantification of trans-synaptic protein alignment: A data analysis case for single-molecule localization microscopy. Methods San Diego Calif 174, 72–80 (2020).

98. Kleinle, J. et al. Transmitter concentration profiles in the synaptic cleft: an analytical model of release and diffusion. Biophys. J. 71, 2413–2426 (1996).

99. Dittrich, M. et al. An excess-calcium-binding-site model predicts neurotransmitter release at the neuromuscular junction. Biophys. J. 104, 2751–2763 (2013).

100. Stiles, J. R., Bartol, T. M., Salpeter, M. M. & Sejnowski, T. J. Synaptic variability: New insights from reconstructions and Monte Carlo simulations with MCell. in *Synapses* 681– 731 (Johns Hopkins University, Baltimore, 2000).

