## Supplementary figures, tables, and methods for "Trans-synaptic molecular context of NMDA receptor nanodomains"

**Contents**

**Figures**

**Supplementary Fig. 1.** Antibody validation

**Supplementary Fig. 2.** DNA-PAINT analysis workflow

**Supplementary Fig. 3.** STED analysis of GluN2 nanocluster relationship

**Supplementary Fig. 4.** Enrichment indices related to Fig. 6

**Supplementary Fig. 5.** Validation of post-fixation surface staining conditions for NMDA treatment experiments, related to Fig. 7

**Tables**

**Supplementary Table 1.** DNA Constructs

**Supplementary Table 2.** Primary antibodies

**Supplementary Table 3.** Secondary reagents

**Supplementary Table 4.** Detailed antibody use in figures

**Supplementary Table 5.** Characteristics of synapses used in modeling

**Supplementary Table 6.** Modeling parameters

**Notes**

**Supplementary Note 1.** Single-molecule analysis pipeline details

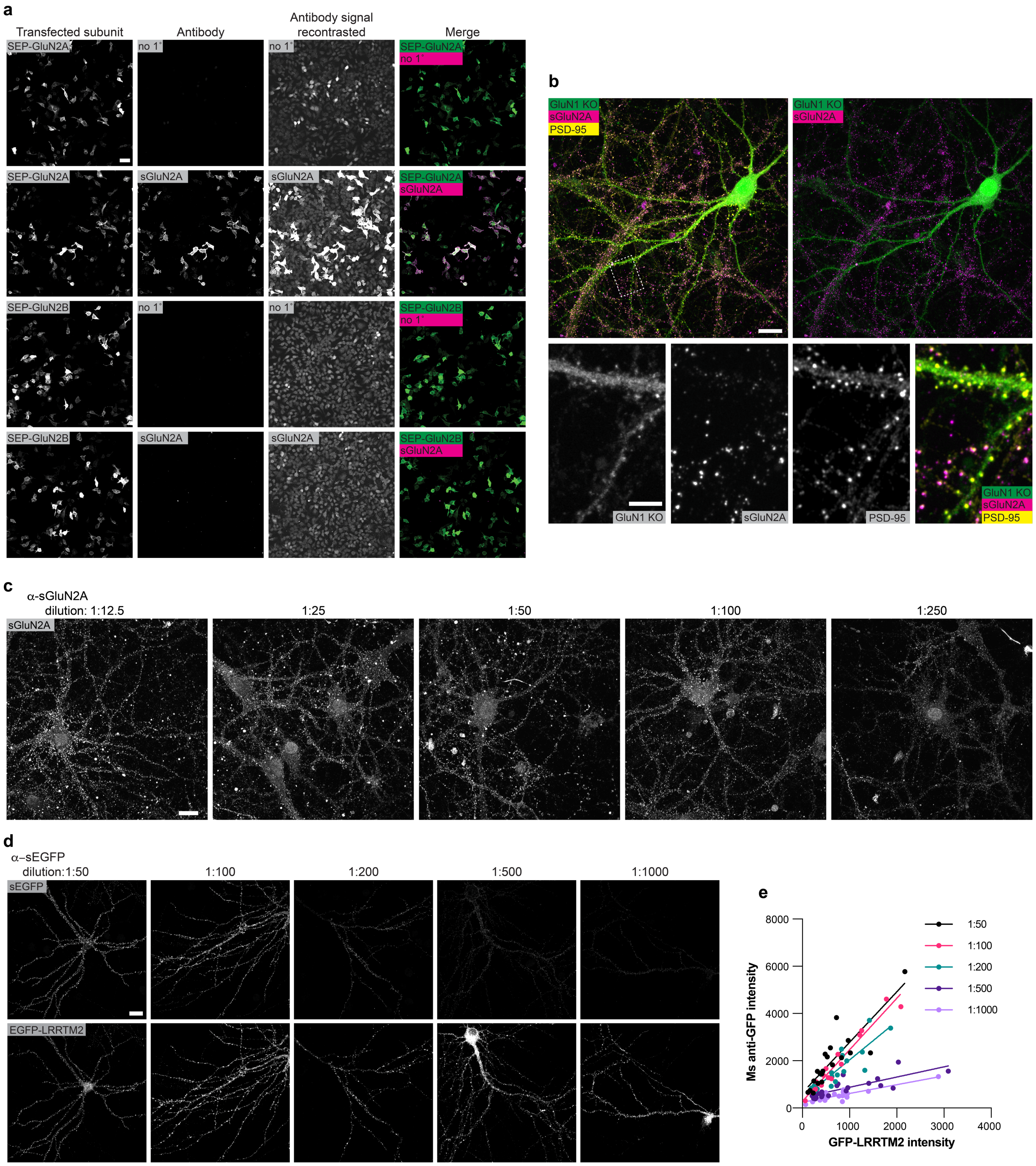


**Supplementary Fig. 1.** Antibody validation

**a** Anti-GluN2A is specific for GluN2A over GluN2B. HEK cells expressing SEP-GluN2A or SEP-GluN2B were surface labeled with anti-GluN2A, and only cells expressing SEP-GluN2A showed anti-GluN2A labeling over background, no primary labeling. Images in the first column are contrast-normalized to the top row image, and in the second column to the second-row image. In the third column, the same images from the second column have been re-contrast-normalized to the image in the first row to show staining background levels. The fourth column is the merge of columns 1 and 2. Scale bar 50 µm. **b** Anti-GluN2A staining is not present in NMDAR knockout neurons. Neurons lacking NMDA receptors due to CRISPR-mediated knockout were not labeled by anti-GluN2A. **(Top)** Images of a neuron expressing EGFP to indicate CRISPR Grin1 KO (green) and stained for surface GluN2A (magenta) and PSD-95 (yellow). Scale bar 20 µm. **(bottom)** Zoom-in of boxed region from **top** where Grin1 KO dendritic spines lacked GluN2A staining but had PSD-95 staining to indicate synapses. GluN2A/PSD-95 positive puncta can be seen on neighboring, wild-type cells in both the zoom and overview images. Scale bar 5 µm. **c** Anti-GluN2A dilution series. Images are contrasted to the 1:12.5 dilution. Large aggregates were apparent in dilutions of 1:50 or higher; thus 1:100 was chosen for experiments in this work. Scale bar 20 µm. **d** Anti-EGFP dilution series. EGFP-LRRTM2 knockdown/rescue-transfected neurons were used as a convenient model system to test a mouse anti-EGFP antibody dilution series to use on EGFP-Grin2b KI cells, as they present a similar synaptic, surface-expressed EGFP epitope, but the number of EGFP-LRRTM2-transfected cells is significantly higher than the number of KIs, allowing for more robust statistics. Neurons were surface stained with mouse anti-EGFP at the indicated dilutions. Images are scaled to the 1:50 dilution for both EGFP and EGFP-LRRTM2. Scale bar 20 µm. **e)** Quantification of total EGFP and anti-EGFP intensity per cell revealed a linear relationship for all dilutions. The saturating 1:50 dilution was chosen for experiments in this work. N = 20, 17, 20, 22, and 18 cells for 1:50, 100, 200, 500 and 1000 dilution, respectively.

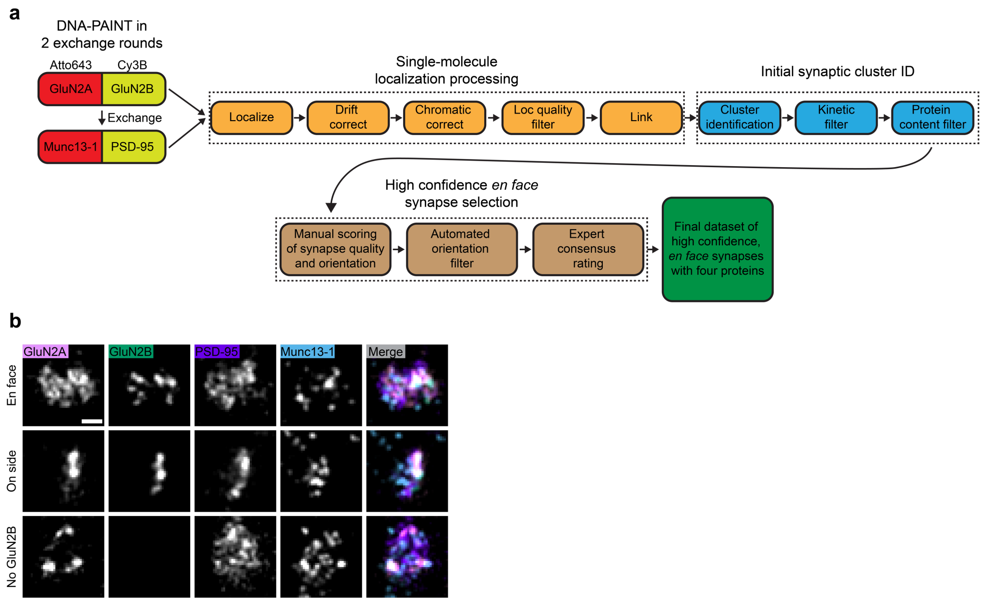


**Supplementary Fig. 2.** DNA-PAINT analysis workflow

**a** Schematic describes four target DNA-PAINT acquisition and data processing workflow. Details can be found in the Supplemental Methods. **b** Example DNA-PAINT renderings (10 nm pixels) show synapses that are oriented perpendicular to the imaging axis (i.e. en face) and used for analysis, or were on their side or lacked GluN2B and discarded from further analysis. Scale bar 100 nm.

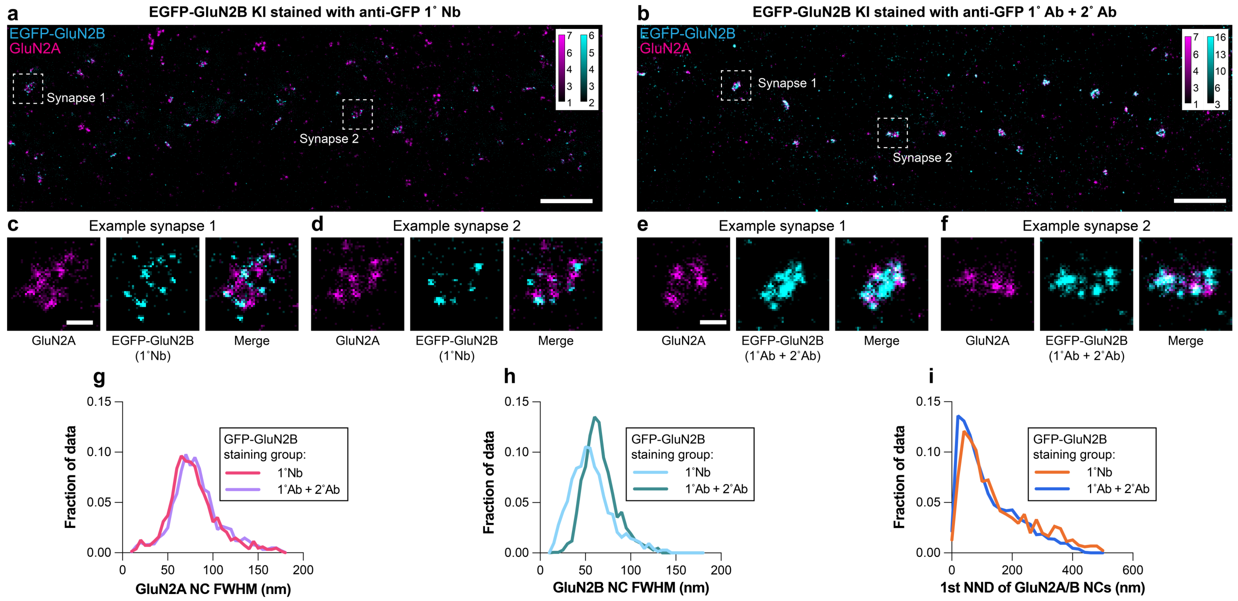


**Supplementary Fig. 3.** STED analysis of GluN2 nanocluster relationship

Antibodies may have limited access to targets in a crowded environment such as the synapse, which could affect interpretation of spatial relationships. To test this, we compared the spatial relationship of GluN2A and GluN2B under favorable (small, direct anti-GFP nanobody) vs unfavorable (large, indirect primary + secondary antibody) steric conditions using STED. These results were consistent with DNA-PAINT and suggest limited effect of sterics on our spatial analysis. **a-b** Exemplar 2D STED images of dendrite stretches from EGFP-Grin2b KI neurons stained for EGFP-GluN2B (cyan) with either (**a**) a direct anti-GFP nanobody (least sterics) or (**b**) a primary plus secondary antibody (worst sterics), along with GluN2A stained with a primary and secondary nanobody in both cases (magenta). Small nanoclusters of both subunits were visually apparent, consistent with our DNA-PAINT results. Intensity scales indicate photon counts. Scale bar 2 µm. **c-f** Boxes indicate zoomed view of synapses in **a-b**. Scale bar 200 nm. **g** GluN2A nanocluster size (average cluster full-width half maximum (FWHM)) is similar between EGFP-GluN2B staining conditions, as expected given GluN2A staining conditions were identical. GluN2A NC N = 1504 for EGFP-GluN2B staining group 1˚Nb, 2043 for staining group 1˚Ab + 2˚Ab. **h** EGFP-GluN2B nanoclusters detected by anti-EGFP nanobody were on average smaller than those detected by primary/secondary antibody, consistent with the smaller linkage error expected for the nanobody vs antibodies. Observing the expected smaller cluster size with primary nanobody is also consistent with our interpretations not being limited by STED resolution. GluN2B NC N = 1403 for EGFP-GluN2B staining group 1˚Nb, 1775 for staining group 1˚Ab + 2˚Ab. **i** The distribution of nearest neighbor distances between GluN2A and EGFP-GluN2B nanocluster peak intensity positions was similar regardless of whether EGFP-GluN2B was detected with a nanobody or antibodies, suggesting our analyses of synaptic spatial relationships were not majorly limited by sterics. N = 1325 distances for staining group 1˚Nb and 1765 for staining group 1˚Ab + 2˚Ab.

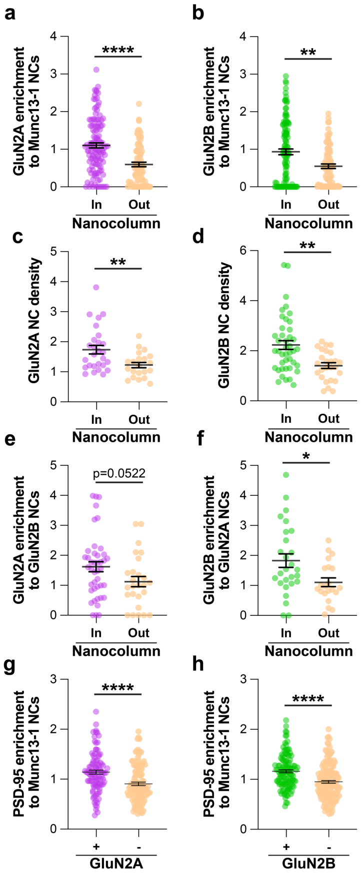


**Supplementary Fig. 4.** Enrichment indices related to Fig. 6

**a-h** Graphs of enrichment indices (average of the first 60 nm of the cross-enrichment plots) from Fig. 6. Panels **a-b** correspond to **Fig. 6b-c**, panels **c-d** correspond to **Fig. 6g-h**, panels **e-f** correspond to **Fig. 6j-k**, and panels **g-h** correspond to **Fig. 6m-n**. N is the same as Fig. 6 per panel. Points are individual nanoclusters of the indicated proteins, lines show mean ± SEM. *p<0.05, **p<0.01, ****p<0.0001.

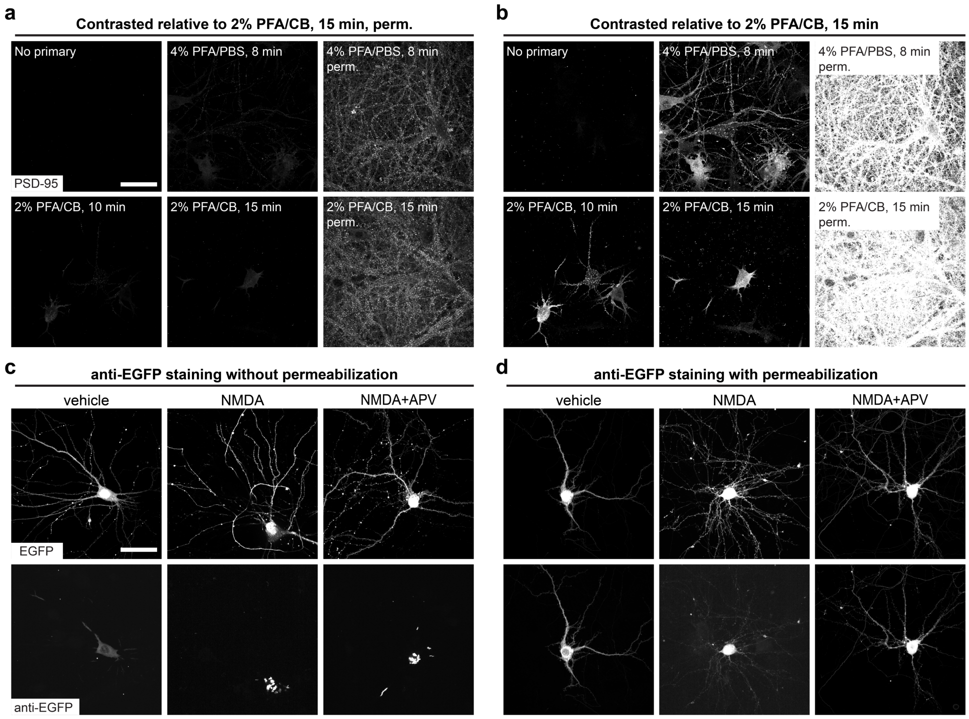


**Supplementary Fig. 5.** Validation of post-fixation surface staining conditions for NMDA treatment experiments, related to Fig. 7

**a-b** 2% PFA/CB fixation for 15 minutes causes only limited permeabilization of neuron dendrites. Neurons in Fig. 7 and Supplementary Fig. 3 were stained for surface epitopes after fixation but before permeabilization. However, fixatives can partially permeabilize cells, which would affect our interpretation of surface receptors. To determine fixation conditions that limited permeabilization, we evaluated detection of an exclusively internal epitope (PSD-95) post-fixation but pre-permeabilization (i.e., surface-stained) with several fixative concentrations, buffers, and timings using post-permeabilization staining as a positive control. Example confocal images show neurons stained for PSD-95 under the conditions indicated. 2% PFA/CB for 15 minutes (used throughout all experiments) resulted in minimal permeabilization as indicated by the general lack of PSD-95 staining, significantly less than 4% PFA/PBS or a shorter 2% PFA/CB fixation. When permeabilization did occur, it was limited to soma or large proximal dendrites, which were easily avoidable for experiments in Fig. 7 and Supplementary Fig. 3. Images in **a** have been contrast-adjusted to match the 2% PFA/CB 15 minutes condition with permeabilization to highlight the low level of post-fixation, pre-permeabilization PSD-95 staining in general. Images in **b** have been contrast-adjusted to match the 2% PFA/CB 15 minutes without permeabilization condition to highlight staining intensities relative to this condition, which was used for experiments. Scale bar 50 µm. **c-d** NMDA treatment does not permeabilize neurons. Neurons in Fig. 7 were treated with drugs that could permeabilize the cell through cytotoxicity, which would affect post-fixation surface staining. To confirm this does not occur, we transfected neurons with cell-fill EGFP and treated with vehicle, NMDA, or NMDA+APV as in Fig. 7, then fixed and stained for EGFP post-fixation, but pre-permeabilization. **c** We observed no significant pre-permeabilization staining of EGFP beyond some soma and large caliber dendrites, consistent with results in **a-b**, and indicating lack of permeabilization by drug treatment. Scale bar 50 µm. **d** By contrast, EGFP staining was apparent in all conditions when staining occurred after permeabilizing the cells, as expected.

**Supplementary Table 1.** DNA Constructs

| Construct | Source | RRID | Figure use |
|---|---|---|---|
| pORANGE GFP-Grin2b KI | Addgene plasmid #131487; http://n2t.net/addgene:131487 | Addgene_131487 | Cloning intermediate |
| pFUGW spCas9 | Addgene plasmid #131506; http://n2t.net/addgene:131506 | Addgene_131506 | 1e-i; 2-3; 5-6; S2b, S3 |
| pFUGW mCherry-KASH | Addgene plasmid #131505; http://n2t.net/addgene:131505 | Addgene_131505 | Cloning intermediate |
| psPAX2 | Addgene plasmid #12260; http://n2t.net/addgene:12260 | Addgene_12260 | Lentivirus production |
| pMD2.G | Addgene plasmid #12259; http://n2t.net/addgene:12259 | Addgene_12259 | Lentivirus production |
| LCV2 | Addgene plasmid # 82416 ; http://n2t.net/addgene:82416 | Addgene_82416 | Cloning intermediate |
| EGFP-LRRTM2 knockdown/rescue | Described in [1] | | S1d |
| SEP-GluN2A | Gift of Andres Barria | | S1a |
| SEP-GluN2B | Gift of Andres Barria | | S1a |
| GluN1-1a pcDNA3.1+ | Gift of Gabriela Popescu | | S1a |
| pFW ORANGE EGFP-Grin2b KI | This manuscript | | 1e-1; 2-3; 5-6; S2b, S3 |
| pFSW myr(Fyn)-EGFP-LDLRct | Described in [2]; specific construct this manuscript | | 1c-d |
| pFW | This manuscript | | Cloning intermediate |
| LCV2 Grin1 KO | Described in [3]; gRNA described in [4] | | S1b |
| pEGFP-N1 | Clontech | | S5 |

**Supplementary Table 2.** Primary antibodies

| Primary antibodies | Source | RRID | Stock | Dilution |
|---|---|---|---|---|
| Monoclonal mouse (IgG2A) anti-GFP (clone 3E6) | Invitrogen A-11120 | AB_221568 | 0.2 mg/ml | 1:50 |
| Polyclonal chicken (IgY) anti-GFP | Invitrogen A-10262 | AB_2534023 | 2 mg/ml | 1:500 |
| Polyclonal rabbit anti-GluN2A | Gift of Rick Huganir; JH6097 | n/a | 0.15 mg/ml | 1:100 |
| Monoclonal mouse (IgG2A) anti-PSD-95 (clone K28/43) | Neuromab 75-028 | AB_2877189 | 0.5 mg/ml | 1:80 |
| Polyclonal rabbit anti-Munc13-1 | Synaptic Systems 126103 | AB_887733 | 0.5 mg/ml | 1:250 |
| Monoclonal mouse IgG2A anti-Bassoon (clone SAP7F407) | Enzo ADI-VAM-PS003 | AB_2313990 | 1 mg/ml | 1:500 |
| GFP-Booster Alexa Fluor 488 | Chromotek gb2AF488 | AB_2827573 | 0.5 mg/ml | 1:500 |
| FluoTag-X2 anti-PSD-95 Atto488 single-domain antibody (sdAb) | Nanotag N3702-At488-L | AB_3076105 | 2.5 µM | 1:500 |

**Supplementary Table 3.** Secondary reagents

| Secondary reagents | Source | RRID | Stock | Dilution |
|---|---|---|---|---|
| FluoTag-XM-QC anti-mouse IgG kappa light chain single-domain antibody (sdAb) (clone 1A23) + docking site F1 | Massive Photonics (custom) | n/a | 5 µM | n/a |
| FluoTag-XM-QC anti-rabbit IgG sdAb (clone 10E10) + docking site F2 | Massive Photonics (custom) | n/a | 5 µM | n/a |
| FluoTag-XM-QC anti-rabbit IgG sdAb (clone 10E10) + docking site F3 | Massive Photonics (custom) | n/a | 5 µM | n/a |
| FluoTag-XM-QC anti-mouse IgG kappa light chain sdAb (clone 1A23) + docking site F4 | Massive Photonics (custom) | n/a | 5 µM | n/a |
| ChromPure mouse IgG, Fc fragment | Jackson Immunoresearch 015-000-008 | AB_2337191 | Variable | n/a |
| ChromPure rabbit IgG, Fc fragment | Jackson Immunoresearch 011-000-008 | AB_2337121 | Variable | n/a |
| Affinipure donkey anti-rabbit IgG | Jackson Immunoresearch 711-005-152 | AB_2340585 | Variable | n/a |

| | | | | |
|---|---|---|---|---|
| Donkey anti-chicken Alexa Fluor 488 | Jackson Immunoresearch 703-545-155 | AB_2340375 | 1.25 mg/ml | 1:500 |
| Donkey anti-mouse (IgG2A) Alexa Fluor 647 | Jackson Immunoresearch 115-605-206 | AB_2338917 | 1.25 mg/ml | 1:500 |
| Donkey anti-rabbit Cy3B | In house | n/a | ~1.25 mg/ml | 1:500 |
| Donkey anti-rabbit Alexa Fluor 647 | Jackson Immunoresearch 711-605-152 | AB_2492288 | 1.25 mg/ml | 1:500 |
| Goat anti-mouse IgG Abberior STAR RED | Abberior STRED-1001-500ug | AB_3068620 | 1 mg/ml | 1:250 |
| FluoTag-X4 anti-GFP Abberior STAR635p sdAb | Nanotag N0304-Ab635p-L | AB_3075902 | 2.5 μM | 1:100 |
| Anti-Rabbit IgG Abberior STAR580 sdAb | Abberior ST580-1010-50UG | Unavailable | 1 mg/ml | 1:200 |
| FluoTag-X2 anti-mouse Ig kappa light chain AZDye568 | Nanotag N1202-AF568 | AB_3075956 | 5 μM | n/a |

**Supplementary Table 4.** Detailed antibody use in figures

| Figures | Antibodies used | Imager used |
|---|---|---|
| 1c-d | (surface) Rb anti-GluN2A + preinc anti-Rb sdAb F2<br>(total) Rb anti-GFP + preinc anti-Ms sdAb F3<br>(total) Ms anti-PSD-95 + preinc anti-Ms sdAb F1<br>(total) Ms anti-Bassoon + preinc anti-Ms sdAb F4 | 1 nM F2-Cy3B (rd 1)<br>0.5 nM F3-Cy3B (rd 1)<br>1 nM F1-Cy3B (rd 2)<br>2 nM F4-Atto643 (rd 2) |
| 1e-f | (surface) GFP-Booster AF488<br>(surface) Rb anti-GluN2A + Dk anti-Rb Cy3B<br>(total) Ms anti PSD-95 + Dk anti-Ms AF647 | |
| 1g-i<br>2-3, 5-6, S2b | (surface) Ms anti-GFP + preinc anti-Ms sdAb F1<br>(surface) Rb anti-GluN2A + preinc anti-Rb sdAb F2<br>(total) Ms anti-PSD-95 + preinc anti-Ms sdAb F4<br>(total) Rb anti-Munc-13 + preinc anti-Rb sdAb F3<br>(total, sequentially after surface antibody staining) GFP-Booster AF488 | 2 nM F1-Cy3B (rd 1)<br>1 nM F2-Atto643 (rd 1)<br>1 nM F4-Atto643 (rd 2)<br>1 nM F3-Cy3B (rd 2) |
| 7 | (surface) Rb anti-GluN2A + preinc anti-Rb sdAb F2<br>(total) Ms anti-PSD-95 + preinc anti-Ms sdAb F4<br>(total) Rb anti-Munc-13 + preinc anti-Rb sdAb F3<br>(total, sequentially after antibody staining) anti-PSD95 Atto488 sdAb | 2 nM F2-Atto655 (rd 1)<br>300 pM F4-Atto655 (rd 2)<br>1 nM F3-Cy3B (rd 2) |
| S1a | (surface) Rb anti-GluN2A + Dk anti-Rb AF647<br>(total) Ck anti GFP + Dk anti-Ck AF488 | |
| S1b | (surface) Rb anti-GluN2A + Dk anti-Rb Cy3B<br>(total) Ms anti-PSD-95 + Dk anti-Ms AF647<br>(total) GFP-Booster AF488 | |
| S1c | (surface) Rb anti-GluN2A + Dk anti-Rb AF647 | |
| S1d | (surface) Ms anti-GFP + Dk anti-Ms AF647<br>(total) GFP-Booster AF488 | |
| S3a | (surface) anti-EGFP STAR635p sdAb<br>(surface) Rb anti-Glun2A + anti-Rb STAR580 sdAb<br>(total, primary sequentially after surface antibody staining) Ck anti-GFP + Dk anti-Ck AF488 | |
| S3b | (surface) Ms anti-GFP + Gt anti-Ms STARRED<br>(surface) Rb anti-GluN2A + anti-Rb STAR580 sdAb<br>(total; primary sequentially after surface staining) Ck anti-GFP + Dk anti-Ck AF488 | |
| S5a-b | Ms anti-PSD-95 + Dk anti-Ms AF647 | |
| S5c-d | Ms anti-GFP + preinc anti-Ms sdAb AZDye568 | |

**Supplementary Table 5.** Characteristics of synapses used in modeling

| Characteristic | Value |
|---|---|
| PSD Area | 0.13 ± 0.02 $\mu m^2$ |
| Number of GluN2A NC per synapse | 5.5 ± 1 |
| Number of GluN2B NC per synapse | 8.1 ± 1 |
| Number of Munc13-1 NC per synapse | 9.4 ± 1 |
| GluN2A NC Area | 844.9 ± 96.8 $nm^2$ |
| GluN2B NC Area | 1040.8 ± 90.9 $nm^2$ |
| Munc13-1 NC Area | 832.0 ± 68.9 $nm^2$ |
| % Munc13-1 NC enriched with GluN2A | 23% |
| % Munc13-1 NC enriched with GluN2B | 29% |
| % Munc13-1 NC enriched with PSD-95 | 24% |

Data are means ± SEM.

**Supplementary Table 6.** Modeling parameters

| Parameter | Value |
|---|---|
| Number of iterations | $1 \times 10^7$ |
| Simulation time step | $1 \times 10^{-7}$ s |
| Output time step | $1 \times 10^{-4}$ s |
| Glutamate diffusion constant | $3 \times 10^{-6}$ $cm^2/s$ |
| Number of glutamate molecules per vesicle | 2200 |
| NMDAR rate constants | Taken from [5] |
| $\tau$ | ~90 μs |
| Synaptic cleft thickness | 15 nm |

**Supplementary Note 1.** Single-molecule analysis pipeline details

1. **Separate emission channels:** Merge multi-page TIFF files from each acquisition in FIJI, then crop the image to separate red and far-red emission and save each as .raw using the ImageJ raw-yaml-export plugin (https://github.com/jungmannlab/imagej-raw-yaml-export).
2. **Localize each emission channel:** Localize spots in each .raw file using the Picasso *localize* command line function (LQ-GPU fit, box size 7, min net gradient set separately for each channel but usually 8000-15000)
3. **Drift correction:** Drift correct images with redundant cross correlation (RCC) using the Picasso *undrift* command line function (segment 1000)
4. **Generate dual view correction t-form:** First roughly align red and far-red sides of the TetraSpeck image by minimizing a nearest neighbor search between bead localizations, then pair localizations between the sides and calculate a t-form using the *fitgeotrans* function in MATLAB with a 2$^{nd}$ degree polynomial. Residual deviation between bead pair positions was estimated as <12 nm after correction.
5. **Recombine acquisitions:** Combine the localization files from each channel of each split acquisition, and correct for the dual view using the t-form and MATLAB's *transformPointsInverse* function to shift localizations into the red emission space. At this point, each region consists of four drift and dual view corrected localization files (for each of four proteins imaged), and operations are performed per region.
6. **Cross-correlate to correct residual offset:** Correct any residual, remaining linear offset between proteins in a region by cross-correlating the images to the GluN2B acquisition. In testing, the gold nanoparticles and overall shape of the image dominated the cross-correlation, and the presence of synapses had little effect. This was performed using a custom MATLAB function based on normalized cross correlation[6], with added image smoothing to remove local high density peaks.
7. **Filter localizations:** For each protein, remove any localizations 1) with fewer photons than the photon mode, 2) standard deviation of the fit >2 pixels (320 nm), 3) localization error >20 nm.

8. **Link localizations:** Link localizations temporally using Picasso *link* command line function with radius 0.3 pixels (48 nm) and 5 dark frames allowed. These parameters were empirically determined to be optimal to link localizations that persisted for more than one frame without linking nearby localizations from another imager strand, and accounting for long-lived localizations that may drop below the detection threshold for several frames.
9. **Cluster identification:** Use Picasso *dbscan* command line function with radius 0.3 pixels (48 nm) and minimum points 10 on each protein in a region to identify densely localized synaptic clusters.
10. **Kinetic cluster filtering:** Using a custom function in MATLAB, filter clusters based on their blinking kinetics[7]. Real DNA imager binding sites will be localized constantly throughout the acquisition, and therefore have a mean frame number close to the total frames/2 (25,000) with a large standard deviation of frame number, while non-specific binding events occur for a few frames then never again, resulting in a low standard deviation of their frame number and a mean frame number that may deviate from total frames/2. Therefore, clusters were removed if their mean frame number was greater or less than the mean +/- 2x the standard deviation of a gaussian fit of the mean frame number of all clusters or if the standard deviation of the frame number was <5000 or >20000 (empirically determined from the images to remove clusters that were not real synapses).
11. **Protein-based cluster filtering:** Using a custom function in MATLAB, PSD-95 clusters were removed from the dataset if they did not have overlap with GluN2B (ie, not a knockin synapse or came from a neighboring cell) or Munc-13 (ie, not a synapse).
12. The result of this process is a drift corrected, dual-view corrected dataset of 4 proteins per region, filtered for high quality localizations, where most non-synaptic and non-KI cell clusters have been removed.
13. **Synapse picking:** To obtain a high-confidence set of near-en-face synapses for analyses:
    a. Each filtered PSD-95 cluster above was manually inspected for the presence of the other imaged proteins in sufficient density for analysis and to confirm that the kept cluster was a synapse. Synapses were kept based on disc like shapes with overlap of pre- and post-synaptic proteins and a size range of ~100 – 800 nm diameter, and their position near a dendrite, then scored as “en face”, “side view” or “intermediate”, ie, somewhere between en face and side.
    b. Synapses were further filtered for en face synapses by removing those from the previously judged “en face” group where the long/short axis ratio of the PSD-95 cluster was > 2.
    c. Finally, 3 expert raters evaluated each synapse as en face and manually adjusted synaptic borders as necessary (only when DBSCAN was obviously incorrect), and in the case of extrasynaptic analysis, manually removed other synapses from within a 500 nm distance of the synapse in question. Synapses were included in the high confidence dataset only when all 3 raters agreed.

For the NMDA treatment experiment, the entire process was essentially identical, but without selecting for GluN2B-containing synapses.

**Supplementary References**


1. Ramsey, A. M. *et al.* Subsynaptic positioning of AMPARs by LRRTM2 controls synaptic strength. *Sci Adv* **7**, eabf3126 (2021).
2. Kameda, H. *et al.* Targeting green fluorescent protein to dendritic membrane in central neurons. *Neurosci Res* **61**, 79–91 (2008).
3. Dharmasri, P. A., DeMarco, E. M., Anderson, M. C., Levy, A. D. & Blanpied, T. A. Loss of postsynaptic NMDARs drives nanoscale reorganization of Munc13-1 and PSD-95. *bioRxiv* 2024.01.12.574705 (2024) doi:10.1101/2024.01.12.574705.
4. Incontro, S., Asensio, C. S., Edwards, R. H. & Nicoll, R. A. Efficient, complete deletion of synaptic proteins using CRISPR. *Neuron* **83**, 1051–1057 (2014).
5. Santucci, D. M. & Raghavachari, S. The effects of NR2 subunit-dependent NMDA receptor kinetics on synaptic transmission and CaMKII activation. *PLoS Comput Biol* **4**, e1000208 (2008).
6. Tang, A.-H. *et al.* A trans-synaptic nanocolumn aligns neurotransmitter release to receptors. *Nature* **536**, 210 (2016).
7. Sun, C. *et al.* The prevalence and specificity of local protein synthesis during neuronal synaptic plasticity. *Sci Adv* **7**, eabj0790 (2021).
